# Brain FGF2 and NCAM1 contribute to FGFR1-dependent progression of ER+ breast cancer brain metastases in young and aged hosts

**DOI:** 10.1101/2025.06.07.658373

**Authors:** Morgan S. Fox, Jenny A. Jaramillo-Gómez, R. Alejandro Marquez-Ortiz, Karen L.F. Alvarez-Eraso, Maria J. Contreras-Zárate, Trinh C. Pham, Elaina N. Barela, Stella N. Koliavas, Peter Kabos, Natalie J. Serkova, Carol A. Sartorius, Elizabeth A. Wellberg, Diana M. Cittelly

**Affiliations:** Department of Pathology; Department of Medicine, Division of Medical Oncology; Department of Radiology, Animal Imaging Shared Resource; University of Colorado Comprehensive Cancer Center, University of Colorado Anschutz, Aurora, CO, 80045, USA; Department of Pathology, Harold Hamm Diabetes Center, and Stephenson Cancer Center, University of Oklahoma Health Sciences Center, Oklahoma City, OK, USA

## Abstract

Estrogen receptor positive (ER+) breast cancer represents a significant proportion of breast cancer brain metastasis but remains understudied. Here, we report that FGFR1-amplification, a well-established driver of ER+ breast cancer endocrine resistance, promotes ER+ breast cancer brain metastatic colonization in young and aged female mice, through both canonical FGF2/FGFR1 signaling and non-canonical NCAM1/FGFR1 interactions. Astrocytic FGF2-mediated paracrine activation of FGFR1 promoted breast cancer brain metastasis in estrogen-treated young mice, but FGF2 levels and signaling decreased in the brain with aging and estrogen-depletion. Neuronal and astrocytic NCAM1, which remain unchanged in young and aged brains, promoted adhesion to neurons, migration, and growth of ER+ cells, suggesting that interactions with astrocytes and neurons facilitate early ER+ breast cancer brain metastasis colonization through FGFR1. Importantly, FDA-approved FGFR inhibitors effectively blocked early colonization but not late-stage brain metastases, suggesting prevention of FGFR1+ brain metastases as a window of opportunity for FGFR1 inhibitors.

## INTRODUCTION

The management of breast cancer brain metastases (BCBM) has emerged as a critical need for metastatic breast cancer (BC) patients^1^. Once diagnosed, brain metastases (BM) remain incurable, and more than 80% of patients die within one year of their diagnosis^2,3^. While BCBM show increased incidence in HER2+ and triple-negative (TNBC, lacking estrogen-receptor (ER), progesterone receptor (PR) and HER2 amplification) subtypes, up to 16% of patients with metastatic ER+ BC develop BM, and represent a large fraction of patients living with BCBM^4–6^.

ER+ BC is effectively treated with anti-estrogen therapies, yet metastatic ER+ BC commonly arises in older women with tumors refractory to these therapies^5^. ER+ BCBM remains understudied, partly due to the dearth of ER+ mouse models that acquire estrogen (17β-estradiol, E2) independence and express drivers of endocrine resistance observed in humans^7–9^. Although human-derived ER+ transplantable tumors and cell lines (patient-derived xenografts, PDXs) are broadly used to study ER+ BC, their growth in mice requires supplementation with E2 mimicking premenopausal levels, as opposed to more relevant post-menopausal or post-endocrine therapy E2 levels in women^7,8,10–12^. Here, we demonstrate that the aged tumor microenvironment (TME) supports the progression of ER+ BC cells in the absence of E2-supplementation, providing clinically relevant models that mimic the natural progression of metastatic ER+ BC.

Mechanisms of acquired and innate resistance to anti-estrogens in ER+ BC have been the focus of extensive research; yet how specific niches at metastatic sites contribute to endocrine resistance and metastatic recurrence are less understood. FGFR1-amplification was recently reported as the only genomic alteration associated with increased risk of late recurrence in post-menopausal women with ER+ early BC on aromatase inhibitor therapy^13^. Aberrations in FGFR signaling have emerged as critical predictors of metastatic progression in ER+ BC following endocrine therapies^13–18^. A recent study showed that ER+ breast tumors acquire an FGFR4-associated signature that correlated with worse prognosis and provided significant and independent prognostic information for brain, liver, and lung metastasis in ER+ tumors only^15^, and suggested that microenvironmental features (either growth factors or specific cell-cell contact) may facilitate growth at metastatic sites. FGFR1 aberrations are significantly higher in BCBM patients compared to non-BM in metastatic BC and predicted poor prognosis^19^, further suggesting that FGFR1 may play a specific role in BCBM promotion.

Tyrosine kinase receptor amplification often results in ligand-independent autoactivation; however, FGFR1-amplification alone is insufficient to drive E2-independent growth in models of ER+ BC primary tumors^20^. In the brain, FGFR1 plays known roles in development and is implicated in synaptic repair and maintenance^21–28^. Although FGFRs typically signal through their cognate FGF ligands, FGFR1 also binds the neural cell adhesion molecule (NCAM), which is abundantly expressed on the surface of neurons and glial cells, and has been shown to activate FGFR1 signaling and regulate synaptic plasticity^21,29–37^. Here, we demonstrate that FGFR1 amplification confers a context-dependent advantage to ER+ BC cells by enhancing brain seeding and outgrowth through microenvironmental activation rather than cell-intrinsic receptor autophosphorylation. ER+ FGFR1+ cells have increased ability to colonize the brain in young and aged/E2-suppressed mice, both through canonical FGF2/FGFR1 signaling and non-canonical NCAM1/FGFR1 signaling. We show that canonical FGF2/FGFR1 signaling decreases in the brain with aging and E2-depletion, and that astrocyte-derived FGF2 drives FGFR1-dependent early brain colonization in young hosts, while NCAM1 serves as a non-canonical FGFR1 activator that promotes brain colonization under aged, low-FGF2 conditions. Importantly, paracrine activation of FGFR1 is critical for early brain colonization but not late-stage brain metastases, suggesting a window of opportunity for FGFR inhibitors to block early colonization in women at high risk of BM.

## RESULTS

### ER+ BCBM models recapitulate clinical progression of ER+ BC metastases throughout the lifespan

Analysis of BCBM samples showed that 41/85 (48.2%) originated from ER+ primary BC, with only 14/41 (34.1%) also harboring HER2-amplification (ER+ HER2+), suggesting that the majority of ER+ BCBM are not driven by HER2-amplification (**Fig. 1a**). Patients with ER+ BCBM were primarily postmenopausal at BM diagnosis, either due to age (>50 years) or prior endocrine therapy (**Fig. 1b,c**). Given the lack of ER+ BC models capable of spontaneous brain colonization, we determined the brain-homing ability of ER+ BC cells following hematogenous dissemination. Four BC PDX-derived cell lines^38^ were injected intracardially in young (<12 weeks) female mice, supplemented with E2 to mimic premenopausal human levels^10^ and required to drive ER+ BC growth in mice^39^. These cells colonized multiple organs, with brain colonization in 0% (UCD46), 20% (UCD4), 40% (UCD12), and 80% (UCD65) (**Fig. 1d**). Since the majority of ER+ BCBM develops following endocrine therapies, we next determined whether ER+ cells could spontaneously disseminate and outgrow upon endocrine therapy/E2-depletion. For this, we implanted UCD4 or UCD65 cells in the mammary fat pad of young ovariectomized (OVX) female NSG mice supplemented with E2 and allowed tumors to grow until reaching 1 cm^3^ to mimic E2-dependent dissemination. Upon tumor removal, mice were randomized into 1) continued E2 supplementation (E2, mimicking E2-dependent metastatic progression in younger women); 2) E2 withdrawal (EWD, mimicking ovarian suppression in younger women); and 3) EWD + letrozole (mimicking full E2-depletion used in post-menopausal women) (**Supplementary Fig. 1a,f**). UCD4 remained E2-dependent for brain and extracranial growth and for dissemination to the brain and other organs (**Supplementary Fig. 1b-e**). BCBM continued growing in a subset of EWD mice, with macrometastasis incidence of 22.2% in the EWD group and 11% in the EWD+LET group. By contrast, E2 did not promote growth of spontaneously disseminated UCD65 cells in the brain but supported growth extracranially (**Supplementary Fig. 1g,h**). While E2 and EWD+LET-treated mice all showed evidence of disseminated UCD65 cells in the brain, mice succumbed to primary tumor recurrence or extracranial metastases (**Supplementary Fig. 1i,j**). Thus, these models fully recapitulate spontaneous ER+ BC metastasis and provide evidence that ER+ BCBM emerge under E2-depletion conditions *in vivo*.

**Fig. 1:**
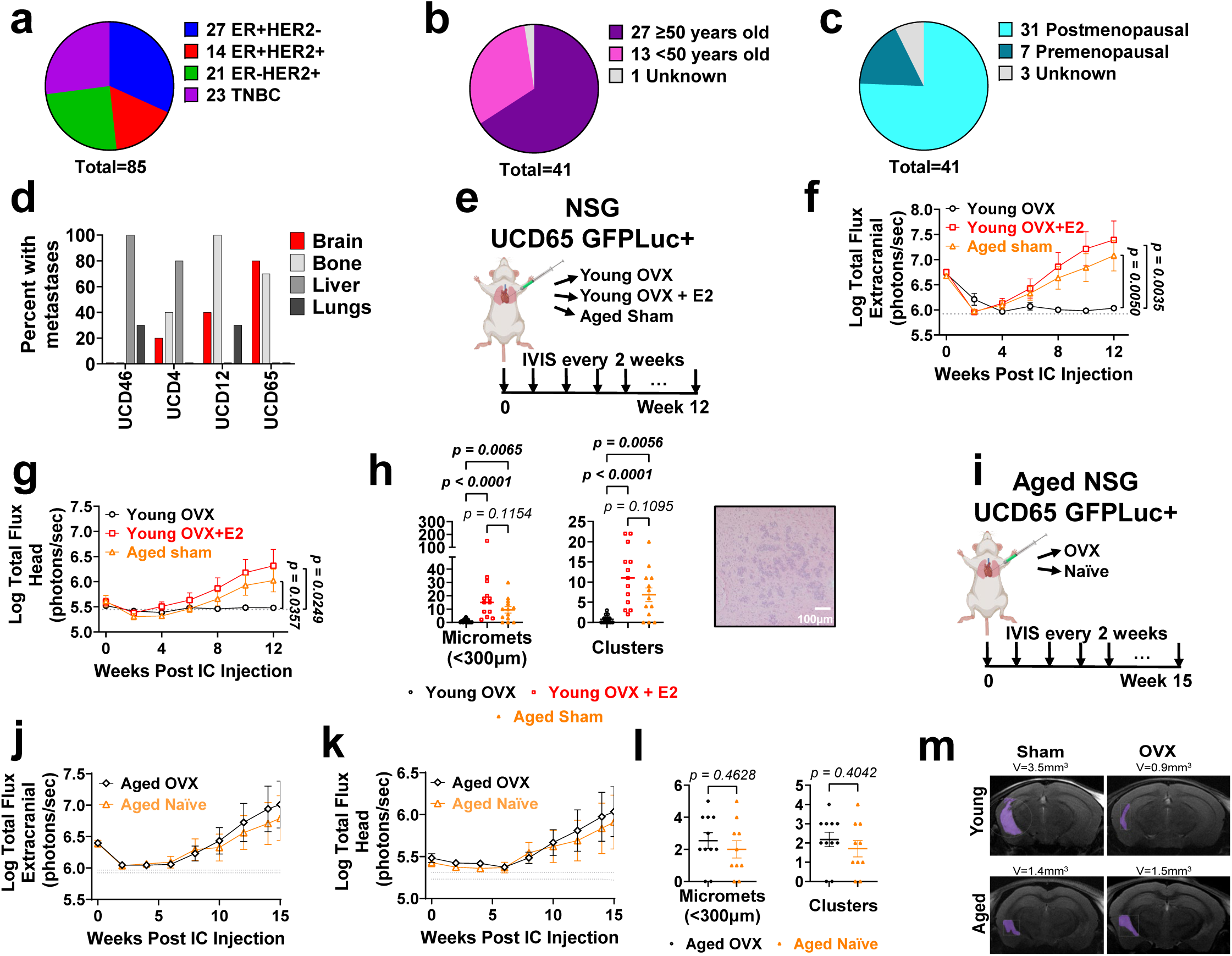
ER+ PDXs effectively model clinical progression of ER+ BC metastases *in vivo*. **a,** A cohort of BCBM from the University of Colorado Cancer Center stratified by primary tumor subtype. **b,** Age distribution of ER+ BCBM patient cohort. **c,** Menopausal status of ER+ BCBM patient cohort. **d,** Frequency of metastasis in brain, bone, liver, and lungs for ER+ PDXs upon intracardiac injection in young mice supplemented with E2 (n=10 per cell line). **e**, UCD65 GFPLuc+ cells were injected intracardially into NSG mice that were i) young OVX (n=14), ii) young OVX+E2 (n=13), or III) aged sham (n=13). **f,** Log-transformed extracranial metastatic burden over time, measured by IVIS. **g,** Log-transformed head metastatic burden over time, measured by IVIS. **h,** Histological quantification BCBM micrometastases and total clusters. Analyzed by Kruskal-Wallis and Dunn’s tests. **i,** UCD65 GFPLuc+ cells were injected intracardially into aged female NSG mice that were i) OVX (n=12) or ii) naïve (n=11). **j,** Log-transformed extracranial metastatic burden over time, measured by IVIS. **k,** Log-transformed head metastatic burden over time, measured by IVIS. **l,** Histological quantification of BCBM micrometastases and clusters. Analyzed by t-test. **m,** High-resolution fast spin echo T2-MRI (axial planes at 9.4 Tesla) of young (12 weeks) and aged (60 weeks) NSG mice that were OVX or naïve, 16 weeks after UCD65 GFPLuc+ intracarotid artery injection, with mask of BM. For **f,g,j,k**, lines denote mean ± SEM. Data were analyzed with 2-way ANOVA or mixed-effects analysis followed by Fisher’s LSD test, with p-value shown for last point only. Gray dotted line denotes baseline IVIS signal from tumor-free mouse.

Given the low BM outgrowth in spontaneous metastases models which prevents mechanistic studies, we used intracardiac injection, mimicking early events of brain colonization, to determine the BM capability of UCD65 cells in young or aged mice (**Fig. 1e**). Consistent with the known E2-dependence of ER+ cells in young mice (<12 weeks), young OVX+ E2 injected with UCD65 cells showed increased extracranial and brain metastasis, with 100% (13/13) mice showing histological evidence of intraparenchymal BM compared to 50% (7/14) in OVX mice. However, in aged mice (>52 weeks), BM were detected in 77% (10/13) and extracranial metastases outgrew in 100% of mice despite low endogenous E2 (**Fig. 1f-h**). BM colonization in aged mice was not dependent on low levels of ovarian E2 as UCD65 cells showed similar progression of BM and extracranial metastases in OVX or naïve aged mice, with histological evidence of intraparenchymal BM in 82% (9/11) and 80% (8/10), respectively (**Fig. 1i-l**). Histological quantification and MRI analysis of young and aged mice harboring UCD65 BMs confirmed the presence of both intraparenchymal micrometastases and macrometastases in both aged groups (**Fig. 1h,m, Supplementary Fig. 1k,l**). Furthermore, histological clusters correlate with head bioluminescence (BLI) signal measured by IVIS (**Supplementary Fig. 1m**). Thus, ER+ BC cells can grow in high E2/young hosts and in low E2/aged hosts.

### FGFR1 drives growth of ER+ BC in the brain TME *ex vivo*

As FGFR1-amplification has been reported as the only genomic alteration that predicts increased risk of late metastatic recurrence in anti-estrogen-treated ER+ BC^13^, we interrogated FGFR1 status in ER+ BCBM clinical samples and cells. Immunohistochemistry showed normal or high FGFR1 expression that did not correlate with membrane pFGFR1 status in ER+ BCBM (**Fig. 2a**). ER+ cells with increased brain-colonization ability (UCD12 and UCD65) harbor genomic amplification and increased FGFR1 mRNA and protein expression compared to the low and non-brain metastatic UCD4, UCD46, and MCF7 cells (**Fig. 2b,c**). FGFR1 amplification in cell lines was not accompanied by increased autophosphorylation (**Fig. 2c**) or expression of cognate FGFR1-activating ligands (**Supplementary Fig. 2a**), suggesting that FGFR1 function may depend on ligand-dependent activation through the TME rather than auto-activation due to high receptor expression. To determine how FGFR1 impacts growth of ER+ BC cells, we stably expressed non-targeting control (shNC) or FGFR1-targeting shRNAs in UCD12 and UCD65 cells (**Fig. 2d, Supplementary Fig. 2b-e**). shFGFR1 and shNC cells showed similar growth in E2-deprived media (vehicle) and in response to E2 in 2D culture (**Fig. 2e,f**). However, when cells were grown as spheroids and plated on organotypic brain slices from young and aged mice, shFGFR1 UCD12 and UCD65 cells showed decreased ability to survive and grow compared to shNC cells (**Fig. 2g-k**). Furthermore, FGFR1-overexpression in MCF7 cells (**Fig. 2l**) did not alter 2D cell growth under E2-deprived or E2-treated conditions compared to empty vector (EV) control cells, but promoted growth in co-culture with organotypic brain slices from young and aged mice (**Fig. 2m-o**). Thus, FGFR1 is a key mediator of survival and growth of ER+ BC cells in the young and aged brain TME.

**Fig. 2:**
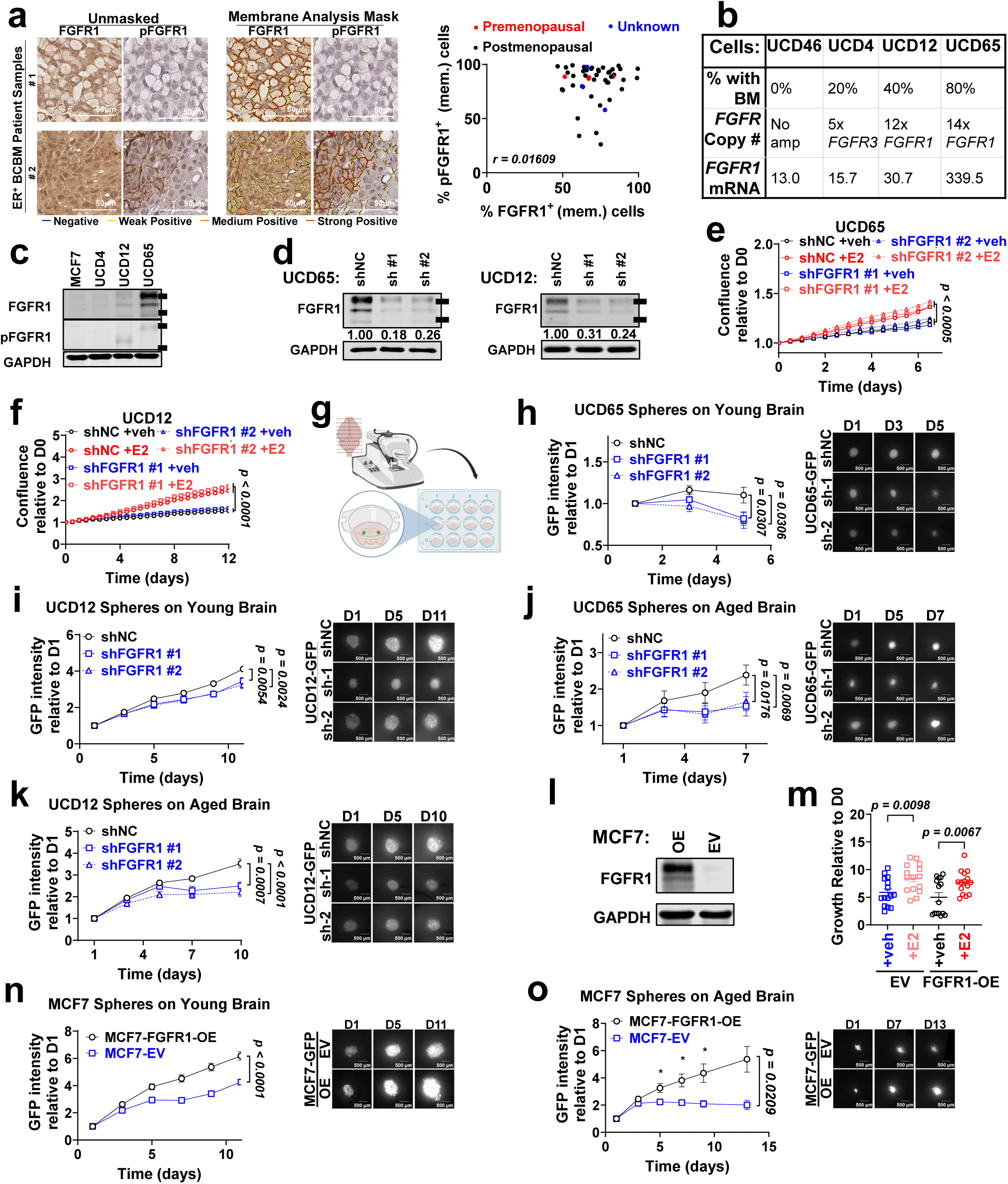
FGFR1 promotes growth of ER+ BC cells in co-culture systems modeling interactions with the brain microenvironment. **a,** IHC of pFGFR1 and FGFR1 in 48 ER⁺ BCBM samples (left) and membrane expression mask (right) performed using the Digital Imaging Aperio algorithm for membrane-expression. Right: Spearman correlation of pFGFR1 and FGFR1 membrane expression. **b,** BM frequency, FGFR amplification status, and FGFR1 relative mRNA (FPKM) for ER+ cells. **c,** FGFR1 and pFGFR1 immunoblots in ER+ BC cell lines. **d,** FGFR1 immunoblot in shNC and shFGFR1 UCD65 and UCD12 cells. Numbers indicate GAPDH-normalized expression relative to shNC, bars indicate 100kDa and 150kDa. Time course growth of shNC and shFGFR1 UCD65 (**e**) and UCD12 cells (**f**, n=10 replicates per condition) in E2-free media, treated with vehicle (EtOH) or 10nM E2. Confluence measured via Incucyte imaging every 12 hours, normalized to time 0. **g,** Schematic of organotypic coculture. UCD65 (**h,** n=12-17) and UCD12 (**i,** n=18-19) spheroid growth on young (9-13w) organotypic brain slices; UCD65 (**j,** n=7-8) and UCD12 (**k,** n=15-16) spheroid growth on aged (49-53w) organotypic brain slices. Graphs show change in GFP intensity per spheroid over time corrected for background fluorescence, normalized to day 1. **l,** FGFR1 immunoblot in MCF7 FGFR1-OE and EV cells. **m,** Fold change growth measured by SRB assay of MCF7 FGFR1-OE and EV cells after 6 days of treatment with 10nM E2 or EtOH vehicle (n=15 replicates per condition). Growth of MCF7 spheroids (n=15-18) on organotypic slices from a young (13w) (**n**) or aged (45w) (**o**) NSG mouse. For **e,f,h-k,n,o**, lines denote mean ± SEM and data were analyzed with 2-way ANOVA followed by Fisher’s LSD test. Data in **m** were analyzed by 1-way ANOVA with Fisher’s LSD test. All scale bars are 500µm. *:p<0.05

### FGFR1 promotes BCBM colonization of ER+ cells in the young and aged brain TME

To test how FGFR1 impacted brain colonization *in vivo,* we intracardially injected shNC or shFGFR1 UCD65 cells into young OVX mice supplemented with E2 (**Fig. 3a**). FGFR1 knockdown (KD) decreased BCBM progression and the number of intraparenchymal metastases but did not affect E2-driven growth of extracranial metastasis (**Fig. 3b-d**). Injection of shNC or shFGFR1 UCD65 in naïve aged mice showed that FGFR1 downregulation decreased BCBM progression, the number of intraparenchymal metastases, and extracranial metastases (**Fig. 3e-h**). Similarly, FGFR1 KD in UCD12 decreased BCBM progression in aged naïve mice and delayed the outgrowth of extracranial metastasis (**Fig. 3i-l**). Moreover, FGFR1-KD cells showed decreased brain signal (but not extracranial signal) immediately following injection (Week 0) in aged mice (**Fig. 3f,j**), suggesting impaired seeding in the aged brain parenchyma. To test if FGFR1 overexpression is sufficient to drive BM colonization and outgrowth, EV or FGFR1-OE MCF7 cells were injected in young/E2 supplemented NSG mice. Despite the poor brain -metastatic ability of MCF7 cells, FGFR1-OE increased BM progression and BM incidence in young OVX + E2 (3/8, 38%) mice compared to OVX- mice (1/8, 12.5%) without impacting E2-driven extracranial metastasis (**Supplementary Fig. 3a-d**). In aged mice, FGFR1-OE cells formed BMs in 14.3% (1/7) mice while EV-MCF-7 cells were unable to colonize (0/8 mice) (**Supplementary Fig. 3e,f**).Together, these data demonstrate that FGFR1 is a key driver of BCBM progression in the young/E2-high TME and promotes brain colonization and extracranial metastases in aged mice.

**Fig. 3:**
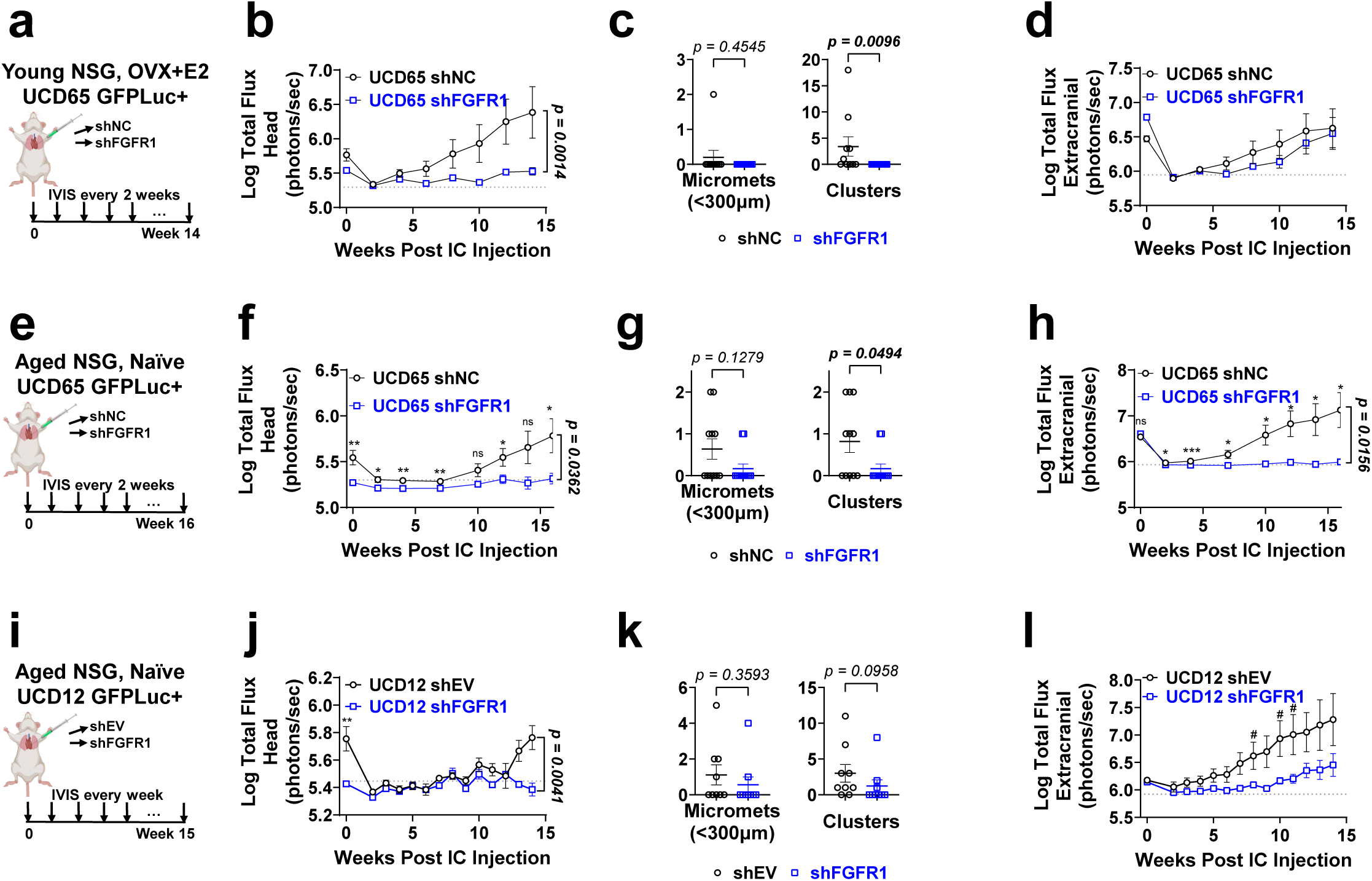
FGFR1 knockdown decreases ER+ BC brain metastasis in multiple models *in vivo*. **a,** shNC (n=12) and shFGFR1 (n=12) UCD65 GFPLuc+ cells were injected intracardially into young (<14w) NSG mice that were OVX+E2. **b,** Log-transformed head metastatic burden over time, measured by IVIS. **c,** Quantification of metastases in brains via H&E. **d,** Log-transformed extracranial metastatic burden over time, measured by IVIS. **e,** shNC and shFGFR1 (n=6-10) UCD65 GFPLuc+ cells were injected intracardially into aged (>52w) naïve NSG mice. **f,** Log-transformed head metastatic burden over time, measured by IVIS. **g,** Quantification of metastases in brains via IHC of GFP and PanCK. **h,** Log-transformed extracranial metastatic burden over time, measured by IVIS. **i,** shEV and shFGFR1 (n=6-9) UCD12 GFPLuc+ cells were injected intracardially into aged (>52w) naïve NSG mice. **j,** Log-transformed head metastatic burden over time, measured by IVIS. **k,** Quantification of metastases in brains via H&E. **l,** Log-transformed extracranial metastatic burden over time, measured by IVIS. For **b,d,f,h,j,l,** lines denote mean ± SEM. Data were analyzed with 2-way ANOVA or mixed-effects analysis followed by Fisher’s LSD test. P-value shown for last time point only. For **c,g,k**, data were analyzed by Mann-Whitney tests. Gray dotted lines denote baseline IVIS signal from a tumor-free mouse. #: p<0.06 *: p<0.05, **: p<0.01, ***:p<0.001.

### FGF2/FGFR1 signaling in glial cells decreases upon E2 suppression and aging

We next interrogated whether host age impacts FGFR1+ ER+ BCBM transcriptional profiles. NanoString GeoMx Digital Spatial Profiling (DSP) followed by UMAP dimension reduction analysis demonstrated no significant differences in the global transcriptional profiles of UCD65 BMs following 6 months of growth in young versus aged sham-mice, suggesting that late-stage ER+ BCBM are transcriptionally similar, independent of host age. Pairwise comparisons revealed only six differentially expressed genes between tumors in young and aged hosts, and Gene Set Enrichment Analysis (GSEA) showed no significant pathway enrichment (**Fig. 4a-e, Supplementary Fig. 4**). To investigate whether changes in the surrounding brain TME due to aging or ovariectomy may impact FGFR1 function in ER+ BCBM, GFAP+ and Iba+ glial areas surrounding PanCK+ UCD65 BMs were analyzed using the mouse whole-transcriptome atlas (**Fig. 4f,g**). Dimension reduction analysis using global transcriptomic profiles showed that glial cells surrounding BMs clustered by age and OVX status, with OVX being a major driver of transcriptional changes and inducing aging-related profiles in young mice (**Fig. 4h**). Pairwise comparisons further demonstrated that E2-depletion and aging broadly repressed gene expression (**Fig. 4i**). To visualize the most representative biological patterns, we performed GSEA and clustered outputs based on shared genes, which identified FGF/FGFR signaling as common top pathways downregulated by OVX and aging (**Fig. 4j-l, Supplementary Data 1**). *Fgf2*, encoding the canonical FGFR1 ligand, was among the top 20 common downregulated genes with OVX and aging (**Fig. 4i,m,n, Supplementary Data 2**), suggesting that paracrine activation of FGFR1 in ER+ BCBM may be driven by glial FGF2 in young hosts, and decreases with aging and E2-depletion.

**Fig. 4:**
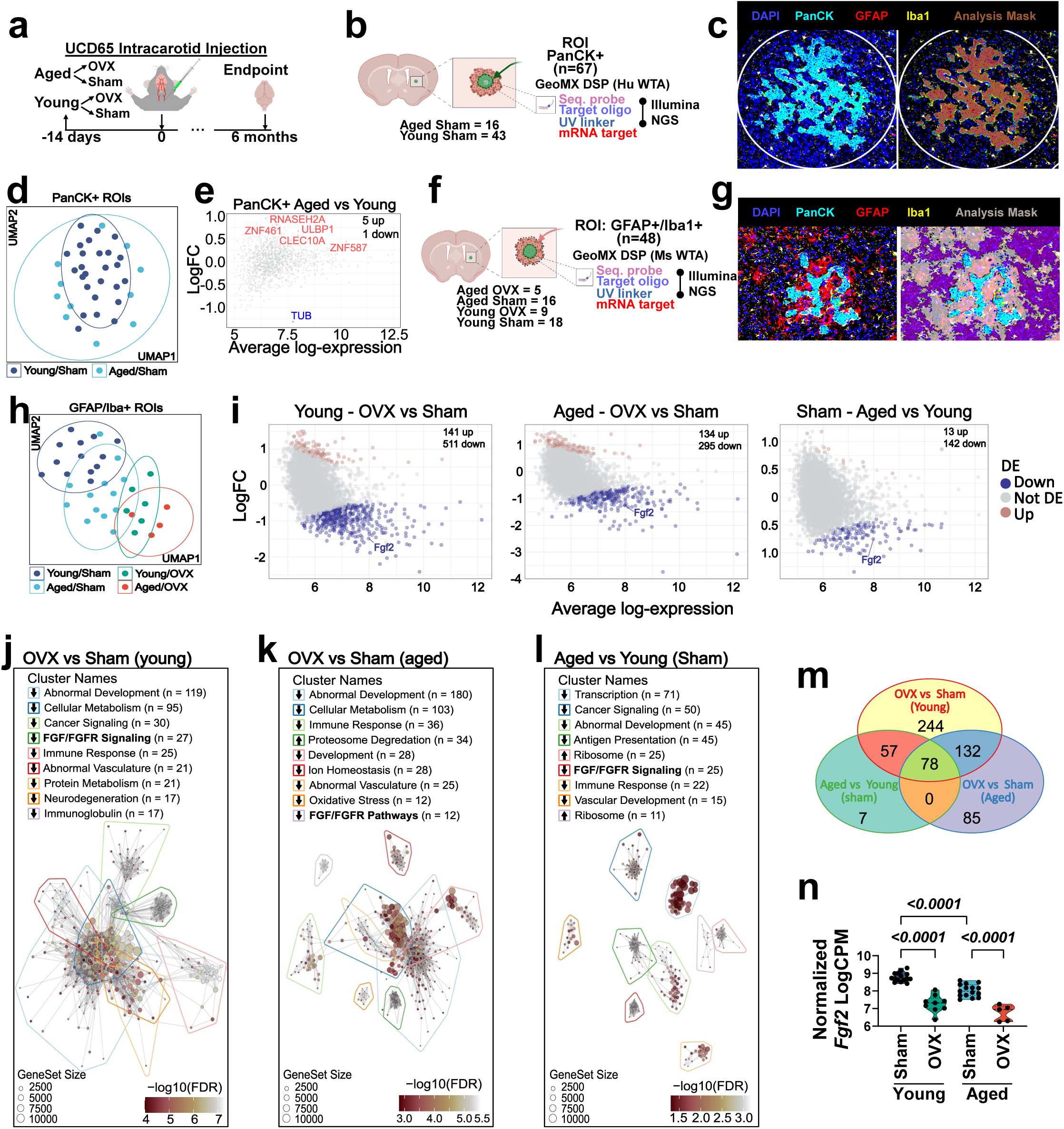
FGF/FGFR signaling decreases in the brain TME with age and E2-depletion. **a,** Aged (<60w) and young (<15w) NSG mice were OVX or sham operated 14 days before intracarotid-injections of UCD65 cells, and brain metastases allowed to progress for six months. At endpoint animals were perfused and brains harvested and sectioned for GeoMX DSP. **b,** mRNA expression in BMs were measured using human whole transcriptome atlas (WTA) library from NanoString in PanCK+ regions of interest (ROI). **c,** Representative image of stained tissue and PanCK+ mask. **d,** Dimension reduction by UMAP of aged and young sham PanCK+ ROIs. **e,** Volcano plots of indicated pairwise comparisons of aged versus young sham PanCK+ ROIs. Significantly differentially upregulated (red) and downregulated (blue) genes are shown. **f,** mRNA expression in glial cells surrounding BMs was measured using mouse WTA NanoString library, using BM-associated GFAP+/Iba1+ ROI. **g,** Representative image of stained tissue and GFAP+/Iba1+ mask. **h,** Dimension reduction by UMAP of GFAP+/Iba1+ ROIs surrounding BM from Young (OVX or sham) and Aged (OVX or sham) mice. **i,** Volcano plots of indicated pairwise comparisons of differentially expressed genes. Significantly differentially upregulated (red) and downregulated (blue) genes are shown. Top 9 differentially expressed clusters of pathways (defined as significant GSEA signatures sharing >35% overlapping genes) for comparisons between young OVX versus sham (**j**), aged OVX versus sham (**k**), and aged versus young sham (**l**) were ordered by number of genesets within the cluster and named based on word frequency in pathway terms. Each node represents a significant geneset, with distance and lines indicating overlap of genes, size representing the number of genes within each geneset, and color representing -logFDR. Repression and enrichment of these gene signatures are indicated with arrows in the legends. **m,** Venn diagram of common downregulated genes in the three indicated pairwise comparisons. **n,** Normalized FGF2 expression within each group (Log(Counts per Million)), analyzed with empirical bayes.

### Astrocytes promote FGF2-dependent activation and proliferation of FGFR1+ ER+ BC cells

We next explored how cells in the brain TME drive canonical FGFR1 activation in young and aged hosts. Consistent with NanoString analysis, FGF2 protein levels were significantly lower in total brain, oligodendrocytes, neurons, and astrocytes from tumor-naïve aged compared to young mice and in astrocytes aged long-term (>3 months) *in vitro* (**Fig. 5a-c**). Since astrocytes expressed the highest FGF2 levels in the brain, we first tested how astrocytic FGF2 impacted FGFR1-dependent ER+ BC growth. UCD65 cells showed increased pFGFR1 when co-cultured with young and aged astrocytes in E2-free media, but activation was higher with young astrocytes (**Fig. 5d**). E2-independent growth of UCD65 cells was higher in co-culture with young astrocytes than with aged astrocytes, and FGFR1 KD significantly decreased growth in co-culture with both young and aged astrocytes (**Fig. 5e-g**). Thus, age-induced changes in astrocytic FGF2 levels correlate with increased FGFR1 activation and growth in ER+ BC, which diminishes with astrocyte age. Similarly, co-culture with young human primary astrocytes promoted UCD12 and UCD65 cell growth and was sensitive to FGFR1 downregulation (**Supplementary Fig. 5a,b),** and blocking FGF2 with a neutralizing antibody decreased UCD12 spheroid growth in organotypic co-culture with young brain slices (**Fig. 5h,i**), suggesting that astrocytic-dependent growth of ER+ BC cells is in part mediated by FGF2/FGFR1 signaling. Consistent with studies showing FGF2 is heparin-bound on the cell membrane rather than secreted^40–42^, FGF2 was not detected in serum-free conditioned media from astrocytes (CM-Ast) but was membrane bound in astrocytes co-cultured with UCD12 cells (**Supplementary Fig. 5c,d**). Moreover, CM-Ast-induced Akt and ERK activation were not altered by BGJ398, a selective FGFR inhibitor that blocks FGF2-mediated signaling and decreases FGF2-dependent proliferation in ER+ BC cells (**Supplementary Fig. 5e,f**). Together, these results suggest that FGFR1 activation by membrane-bound FGF2 is critical for tumor proliferation in the absence of exogenous E2, and that young but not aged astrocytes can robustly support E2-independent proliferation of FGFR1+ BC cells.

**Fig. 5:**
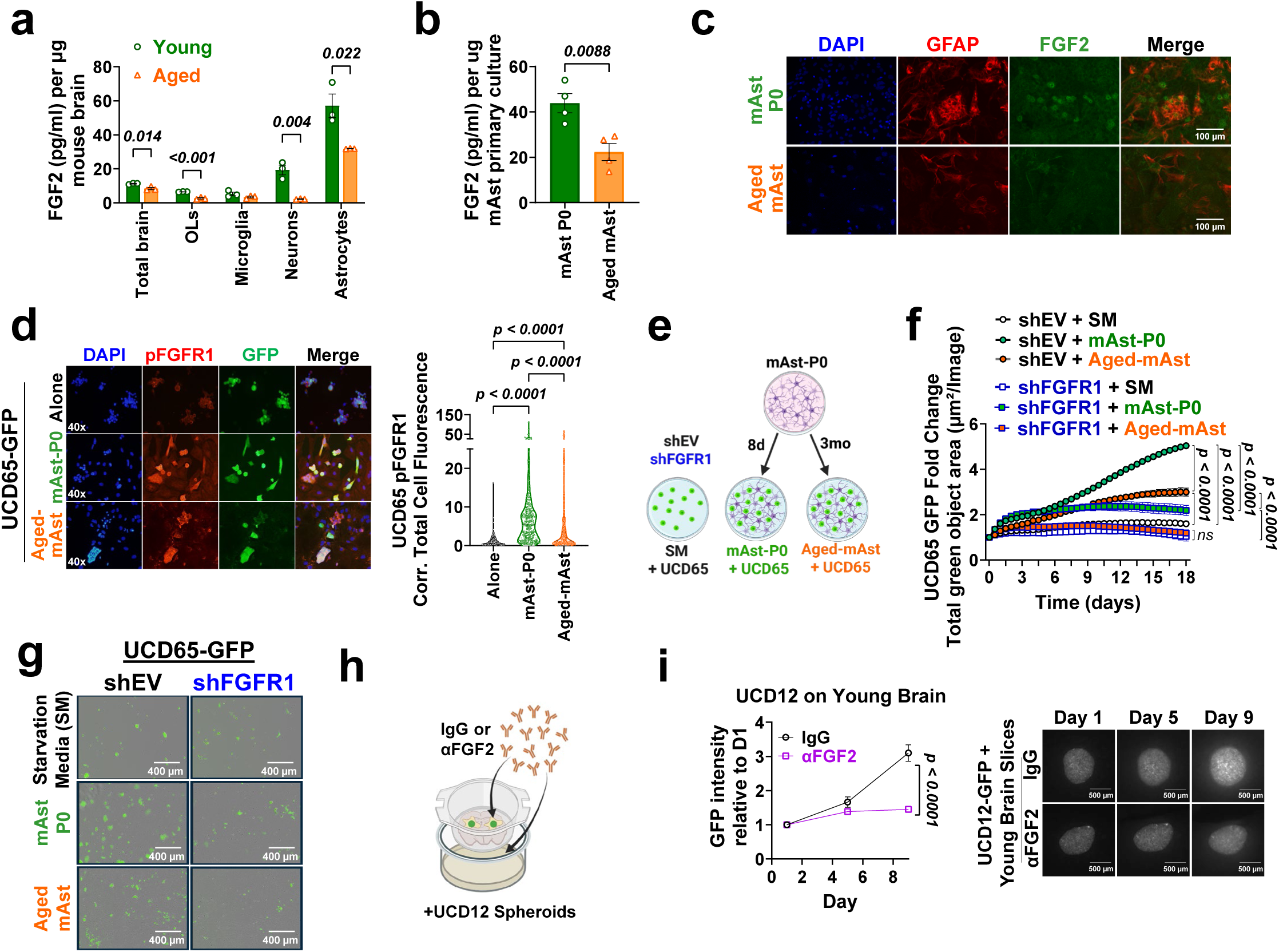
Astrocytes promote canonical FGF2-dependent activation of FGFR1 in ER+ BC. **a,** FGF2 ELISA from total brain, oligodendrocyte (OL), microglia, neuron, and astrocyte lysates isolated from young (12 weeks) and aged (>56 weeks) female mice (n=3). **b,** FGF2 ELISA from young (mAst P0, 15 days *in vitro*) and aged (Aged mAst, 90 days *in vitro*) mouse primary astrocyte lysates (n=4). **c,** Immunofluorescence of GFAP and FGF2 in mAst-P0 and aged-mAst. **d,** Immunofluorescence of pFGFR1 in UCD65 alone or co-cultured with mAst-P0 and aged-mAst in serum-free media. Right: pFGFR1 Corrected Total Cell Fluorescence in UCD65 (n=228-303 cells per group). **e**, Schematic of the experimental design for co-cultures of UCD65 shEV and shFGFR1 cells with mAst-P0 or aged-mAst. **f**, Fold change in total green area relative to time 0 of UCD65 shEV and shFGFR1 cells cultured alone or with mAst-P0 or Aged-mAst in E2-free media (n=3 replicates per condition), measured by Incucyte. **g**, Representative images at 18 days. **h,** Schematic of organotypic co-cultures with neutralizing antibodies. **i,** Spheroids of GFP+ UCD12 cells (n=10 per condition) were plated on organotypic brain slices from a young (8w) NSG mouse. Slices were pretreated for 24h with 1µg anti-FGF2 blocking antibody or IgG control. Graph shows change in GFP intensity per spheroid over time corrected for background fluorescence, normalized to day 1. For **a,b,** data were analyzed with t tests. **d** was analyzed with Kruskal-Wallis test followed by Dunn’s multiple comparison test. Adjusted P-value shown. For **f,i** line denotes mean ±SEM, data were analyzed with 2-way ANOVA followed by Fisher’s LSD test. P-values shown for last time point only.

### NCAM1 and FGF2 activate FGFR1 signaling and growth in ER+ BC

Given that brain FGF2 levels decrease with age and E2-suppression, and aged astrocytes were less effective in promoting FGFR1-dependent growth, we investigated whether non-canonical mechanisms of FGFR1 function could impact ER+ BCBM in the absence of high FGF2. NCAM1 is a neuronal and astrocytic cell adhesion molecule whose expression is unchanged with aging (**Supplementary Fig. 6a- d**) and is known to activate FGFR1 and regulate neural development and synaptic plasticity^31,37,43–45^. To determine if NCAM1 could activate FGFR1 kinase signaling in ER+ BC cells, we treated UCD65 cells with NCAM1 and found dose-dependent activation of FGFR1, Akt, and ERK (**Fig. 6a**), even at the lowest dose (which is below reported NCAM1 cell surfaces levels in the brain^31,32^). NCAM1-induced signaling is dependent on FGFR1 expression, as it was decreased in shFGFR1 compared to shNC UCD65, and only observed in MCF7-FGFR1-OE but not MCF7-EV cells (**Fig. 6b,c**). BGJ398 blocked FGF2 and NCAM1-induced FGFR1 signaling in UCD65, UCD12, and MCF7-FGFR1-OE at multiple time points (**Fig. 6d-f, Supplementary Fig. 6e,f**). Since NCAM1 is mostly membrane-anchored, we next tested whether surface NCAM1 from neighboring cells activates FGFR1 by co-culturing ER+ BC cells with NCAM1+ SY5Y neuroblastoma cells stably expressing shEV or shNCAM1. Co-culture with shNCAM1 SY5Y cells significantly reduced FGFR1 phosphorylation in all ER+ cell lines compared to shEV controls (**Fig. 6g,h, Supplementary Fig. 6g,h**). Similarly, co-culture with astrocytes transduced with mCherry-tagged AAV expressing shNCAM1 reduced pFGFR1 in UCD65 and UCD12 cells compared to EV control, demonstrating that NCAM1 from both neurons and astrocytes can activate FGFR1 in ER+ BC cells (**Fig 6i,j, Supplementary Fig. 6i**). A blocking antibody against NCAM1 decreased UCD12 spheroid growth in aged, but not young organotypic brain slices, suggesting that NCAM1 promotes growth of ER+ BC in the aged/low FGF2 brain TME, but it is not the main driver in the high FGF2 young TME (**Fig. 6k,l**). Because young astrocytes express both NCAM1 and FGF2 at high levels, we next examined their relative contribution to ER+ BC proliferation. Lentiviral-mediated KD of NCAM1 and FGF2 effectively decreased FGF2 and NCAM1 levels in astrocytes, resulting in significant decreased proliferation of UCD12 and UCD65 cells compared to co-culture with shNC astrocytes. Astrocytic FGF2 KD produced a greater reduction in ER+ BC growth than NCAM1 KD, with minimal additive effects upon combined depletion of NCAM1 and FGF2 (**Fig. 6m,n, Supplementary Fig. 6j,k**). Thus, these results indicate that while astrocyte-derived FGF2 remains the dominant driver of FGFR1-dependent growth in young brains, NCAM1 serves as a non-canonical FGFR1 activator that sustains ER⁺ BC growth under aged, low-FGF2 conditions.

**Fig. 6:**
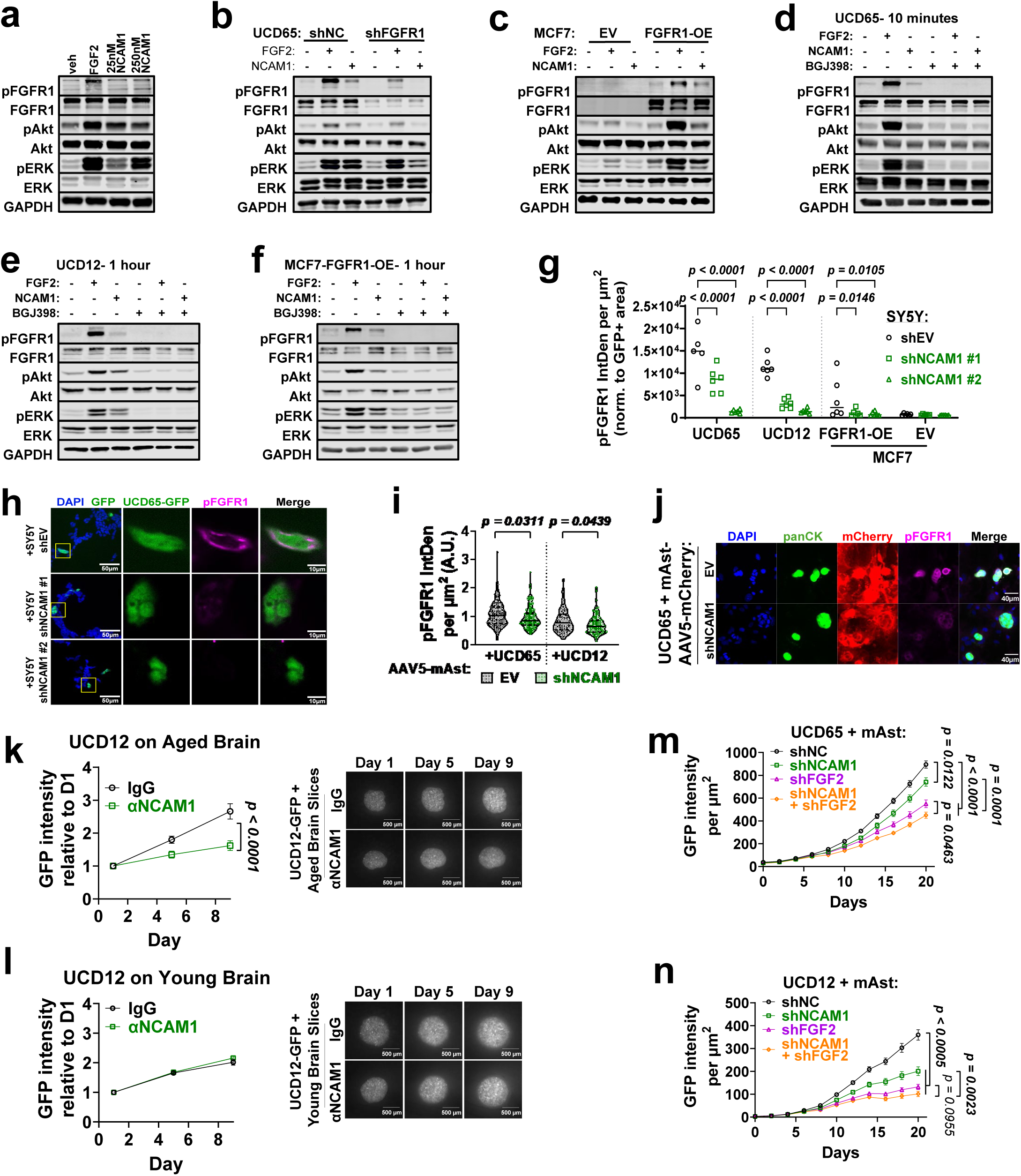
NCAM1 and FGF2 activate FGFR1 signaling and growth in ER+ BC. **a,** UCD65 cells were serum-starved and then treated with FGF2 (10ng/mL), 25nM or 250nM recombinant human NCAM1, or vehicle for 10 minutes. Immunoblot shows pFGFR1, FGFR1, pAkt, Akt, pErk, ERK. GAPDH is loading control. **b,** Immunoblot from UCD65 shNC and shFGFR1 cells treated with 10ng/mL FGF2, 250nM NCAM1, or vehicle for 1 hour. **c,** Immunoblot from MCF7 EV and FGFR1-OE cells treated with 10ng/mL FGF2, 250nM NCAM1, or vehicle for 1 hour. **d-f**, Immunoblot from UCD65, UCD12, and MCF7 FGFR1-OE cells pretreated with 10µM BGJ398 or vehicle for 1 hour, then treated with 10ng/mL FGF2, 250nM NCAM1, or vehicle for the indicated times. **g,** Quantification of pFGFR1 immunostaining in UCD65, UCD12, MCF7-FGFR1-OE, and MCF7-EV cells cultured on top of shEV or shNCAM1 SH-SY5Y cells in serum-free media (n=5 replicates per condition). Graph shows pFGFR1 Intensity in GFP+ areas. **h,** Representative images of UCD65 cells co-cultured with shEV and shNCAM1 SH-SY5Y cells. **i,** UCD65 and UCD12 cells were grown for 7 days in starvation media on coverslips in co-culture with AAV5-mCherry-mAsts. IF of panCK and pFGFR1. pFGFR1 intensity was quantified within panCK+ ROIs (n=108-129 for UCD65, n=60-89 for UCD12). **j,** Representative images of UCD65-mAst co-cultures. Spheroids of GFP+ UCD12 cells (n=10 per condition) were plated on organotypic brain slices from aged (46w, **k**) or young (8w, **l**) NSG mice. Slices were pretreated for 24h with 1µg anti-NCAM1 blocking antibody or IgG control. Graph shows change in GFP intensity per spheroid over time corrected for background fluorescence, normalized to day 1. GFP+ UCD65 (**m**) and UCD12 (**n**) cells were co-cultured on transuced mAsts (n=6 per condition) for 20 days in starvation media. GFP intensity was measured every two days via Incucyte. **g,i** were analyzed by one-way ANOVA followed by Fisher’s LSD test. **k-n** were analyzed by two-way ANOVA followed by Fisher’s LSD test. p-values shown for final time points.

### NCAM1/FGFR1 communication drives interactions between ER+ BC and neurons

Studies have shown that FGF2 and NCAM1 activation of FGFR1 can result in different downstream cellular effects^44^. To define downstream mechanisms resulting from NCAM1 or FGF2-mediated FGFR1 activation in ER+ BC cells, we performed global gene expression profiling of UCD12 and UCD65 cells treated with vehicle, NCAM1, or FGF2 for 24h (**Fig. 7a, Supplementary Fig. 7a-c**). GSEA identified gene signatures shared by both cell lines that were commonly enriched or repressed by NCAM1 and FGF2 or uniquely regulated by each ligand (**Fig. 7b, Supplementary Data 3**). FGF2 and NCAM1 commonly induced 1,390 gene signatures, primarily clustered as signaling associated with development and disease, DNA replication, and immunity. FGF2 induced 2,879 unique differential signatures, demonstrating enrichment of canonical growth factor receptor signaling, cytoskeleton organization, ECM interactions, and transmembrane transport and secretion (**Supplementary Fig. 7d,e, Supplementary Data 4,5**). By contrast, NCAM1 treatment induced 90 unique differential gene signatures, primarily classified by disease and cell signaling activation, metabolic changes, transmembrane transport, and, interestingly, neuronal gene signatures (**Fig. 7c, Supplementary Fig. 7f,g**). These neural signatures showed enrichment in differentiation, development, and adhesion, with repression of postsynaptic membrane protein localization and trafficking, and myelination/ensheathment (**Fig. 7d**). Therefore, these results suggest that FGF2 promotes broad transcriptional changes, driving canonical signaling through FGFR1 and proliferation, while NCAM1 may drive specific gene signatures promoting neural interactions in ER+ BC cells.

**Fig. 7:**
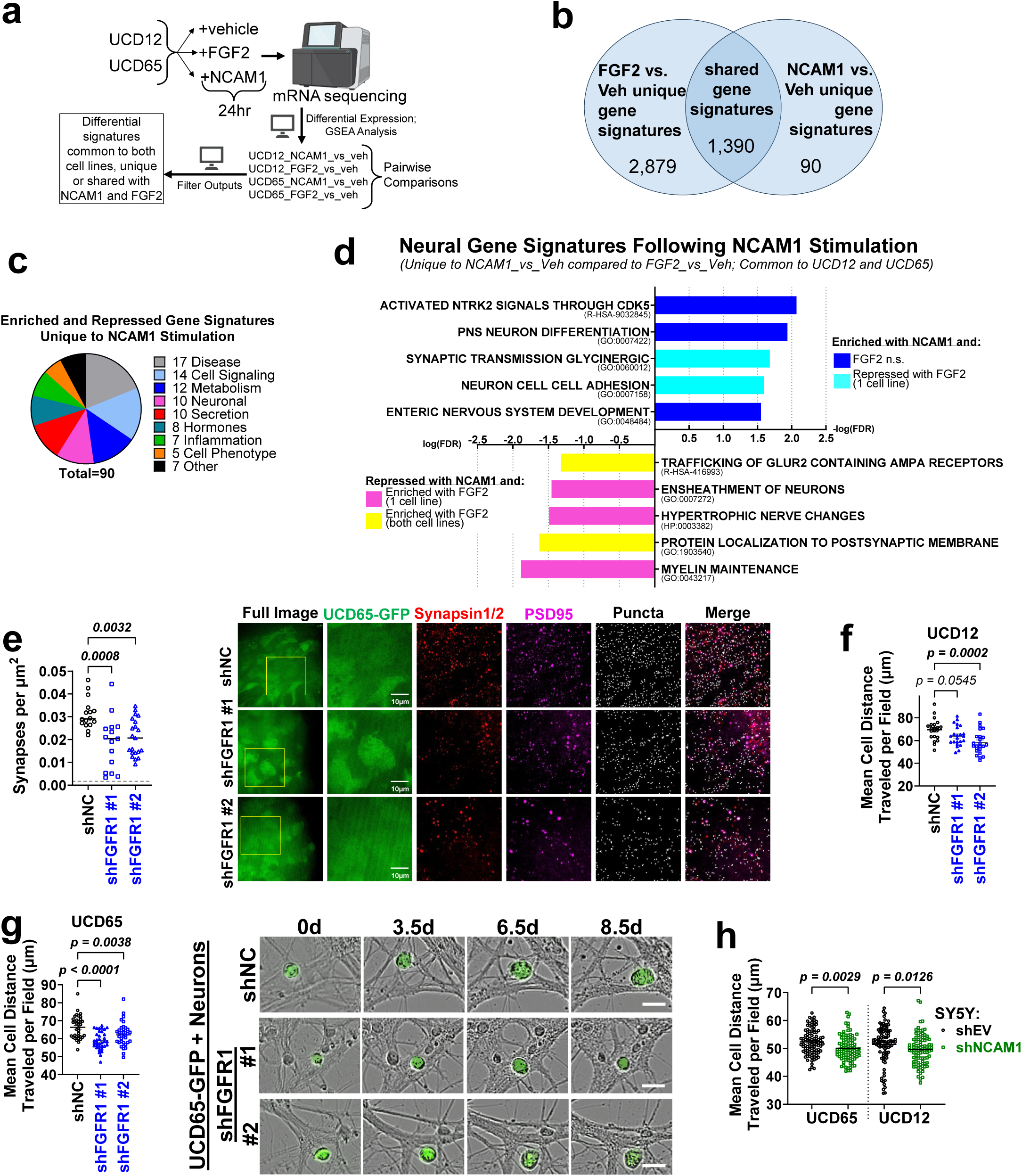
NCAM1/FGFR1 drives interactions between ER+ BC and neurons. **a,** RNA-sequencing schematic for UCD12 and UCD65 cells in serum-free media treated with vehicle, 10ng/mL FGF2, or 20ug/mL NCAM1 for 24hr (n=3). **b,** Venn diagram shows unique and shared significant gene signatures following FGF2 and NCAM1 treatment. **c,** Pie chart of gene signature categories differentially induced by NCAM1 treatment. **d,** log(FDR) of significant neural gene signatures differentially enriched or repressed following NCAM1 treatment. **e,** Immunofluorescent staining of PSD95 and Synapsin1/2 in shNC and shFGFR1 UCD65 spheroids in co-culture with organotypic brain slices from aged NSG female mice (>59w) for 3 days. Punctate colocalization (n=15-21 images) was quantified. 5,000 GFP+ shNC and shFGFR1 UCD12 (**f**) or UCD65 (**g**) cells were plated on coverslips with 315,000 primary hippocampal neurons (DIV: 8) and imaged every 12 hours for 6-9 days via Incucyte. Cell migration was measured and mean distance per cell within each image (n=21-36) was plotted. Right: representative images of UCD65 cells. Scale bars: 20µm. **h**, 5,000 UCD12 and UCD65 cells were plated on a monolayer of shEV or shNCAM1 SY5Y cells and imaged every 12 hours for 6.5 days via Incucyte. Cell migration over time was measured and mean distance per cell within each image (n=96) was plotted. For **e-h**, data were analyzed by 1-way ANOVA followed by Fisher’s LSD test.

Given the known roles of NCAM/FGFR1 interactions in modulating synaptic plasticity, the induction of synaptic-related gene signatures upon NCAM1 treatment in UCD65 and UCD12 cells, and emerging evidence that direct and indirect synaptic communication between neurons and disseminated brain metastatic cells are key mediators of BCBM outgrowth^46–48^, we next determined how FGFR1 impacts cancer-neuron interactions. FGFR1 KD resulted in decreased density of excitatory synapses at the interface of UCD65 spheroids and organotypic brain slices, suggesting that FGFR1 contributes to synaptic localization or interaction between ER+ BC cells and organotypic brain slices (**Fig. 7e**). Furthermore, coculture with primary hippocampal neurons from postnatal mice demonstrated that ER+ BC cells bind and migrate along neurites *in vitro*, and FGFR1 KD significantly decreased the distance migrated by UCD65 and UCD12 cells along these neurites (**Fig. 7f,g**). NCAM1 KD in SY5Y cells decreased migration of UCD65 and UCD12 cells in co-culture (**Fig. 7h**). Together, these results suggest that FGFR1 promotes neuronal interactions and that FGFR1/NCAM1 facilitates migration of ER+ BC cells.

### FGFR1 kinase inhibition is effective only in the prevention of ER+ BCBM

Our data suggests that paracrine activation of FGFR1 by FGF2 and NCAM1 can influence early colonization as well as late metastatic progression of ER+ BC. Since interactions with astrocytes and neurons occur from early stages of metastatic colonization, we assessed UCD65 FGFR1 activation 5 days following intracardiac injection in young and aged hosts. Membrane pFGFR1 expression was found in 83% of early disseminated cells in young mice and 57% in aged mice, and membrane pFGFR1 intensity was stronger in disseminated cells in young compared to aged brains (**Fig. 8a,b**). These findings are consistent with strong FGF2-dependent FGFR1 activation of disseminated cancer cells in young mice, and weaker, NCAM-dependent activation in aged mice. In contrast, late-stage UCD65 BCBM from young and aged mice showed minimal membrane pFGFR1 (**Fig. 8c**), paralleling the lack of correlation between pFGFR1 status and total FGFR1 expression observed in late-stage clinical samples (**Fig. 2a**). Thus, this data suggests that paracrine activation of FGFR1 is prominent during early brain colonization, but it is dispensable during later stages of BM progression. To test this, we injected UCD65 cells in young mice supplemented with E2 or in aged naïve mice (**Fig. 8d,i**) and began treatment with vehicle or BGJ398 either immediately after injection or upon detectable brain metastatic outgrowth. We confirmed the reported brain-penetrance of BGJ398^49^, as it effectively blocked TME-mediated FGFR1 activation in disseminated cancer cells at the dose used (**Supplementary Fig. 8a**). In young mice, early FGFR inhibition significantly suppressed BCBM progression, reducing IVIS-detectable BM as well as the number of micrometastases and BM clusters when treatment was initiated at the time of cell injection, but not when initiated 7 weeks post-injection (**Fig. 8e-h).** Consistent with the FGFR1 independence of advanced lesions, IF analysis revealed no significant differences in pERK staining in late-stage BMs following BGJ398 treatment (**Supplementary Fig. 8c,d**). In aged mice, characterized by weaker FGFR1 activation and slower BM progression, BGJ398 treatment did not significantly affect BM progression or *ex vivo* IVIS (**Fig. 8j,k**), nor did it affect extracranial progression (**Supplementary Fig. 8e**). Although BGJ398-associated toxicity in aged mice required the termination of these studies before control mice reached euthanasia criteria, histological quantification of BMs revealed that early treatment BGJ398 reduced the total number of metastatic clusters and micrometastases (**Fig. 8l**). These findings support a role for FGFR1 activation in early brain colonization, even in the aged brain, while suggesting that established metastases are largely FGFR1-independent.

**Fig. 8:**
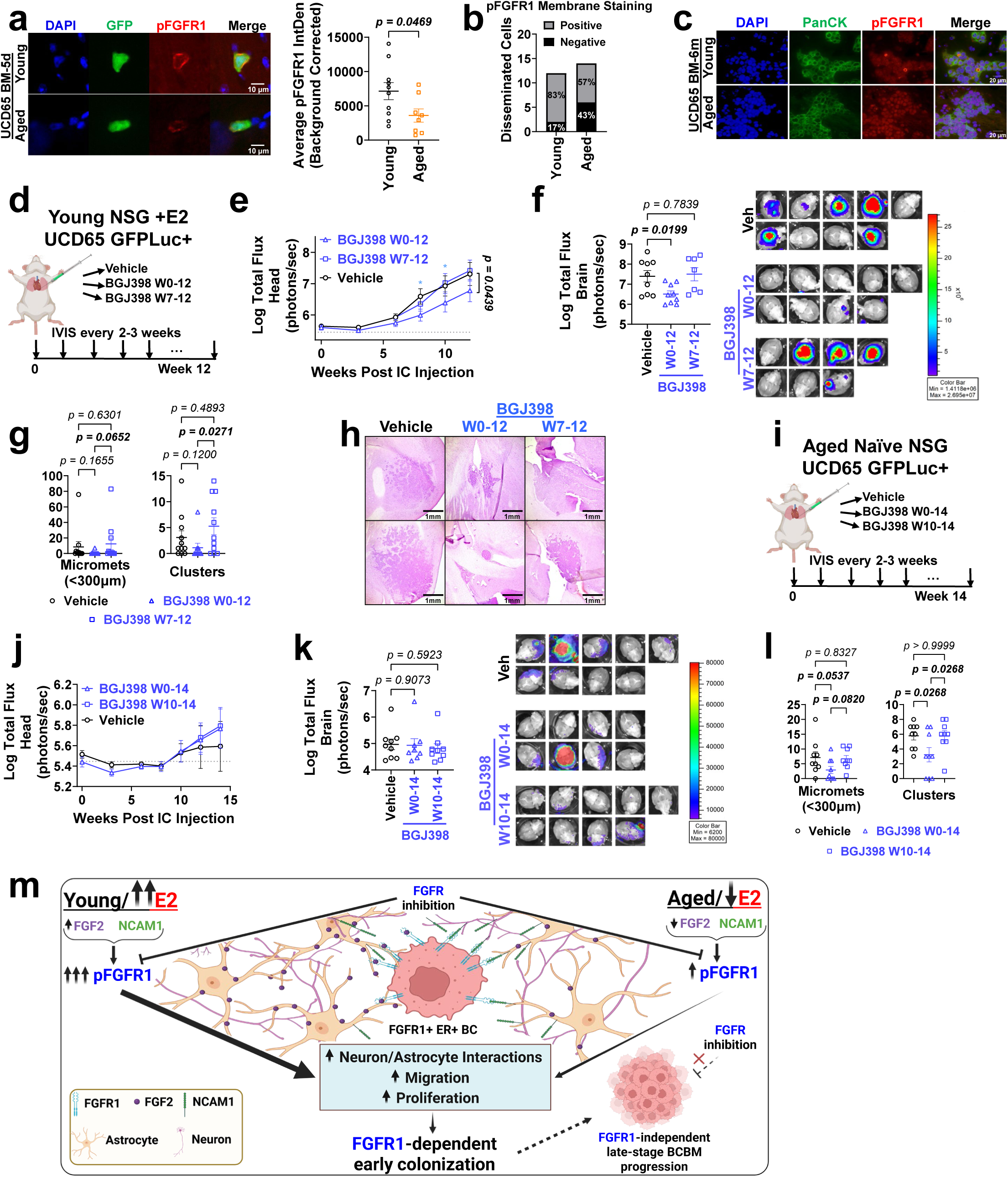
FGFR1 kinase inhibition is only effective in early BCBM. **a,** pFGFR1 quantification in brain disseminated UCD65 cells 5 days after intracardiac injection in young (<12w) and aged (>56w) NSG mice. Graph shows average pFGFR1 intensity in GFP+ ROIs (n=8-10 images per group) corrected by background (GFP-). Data analyzed by Welch’s t-test. **b,** number and percentage of brain disseminated cells with positive membrane pFGFR1 staining. **c,** IF staining of PanCK and pFGFR1 in brain metastases from UCD65 cells in young (<15w) and aged (>60w) NSG mice, 6 months post-intracarotid injection. **d,** UCD65 GFPLuc+ cells were injected intracardially into OVX young (<18w) NSG mice treated with E2 and randomized to receive vehicle (1% CMC, n=11) or BGJ398 (30mg/kg) 5x/week PO starting at the time of injection (W0-12, n=9) or 7 weeks post-ic (W7-12, n=11). **e,** Log-transformed head metastatic burden over time measured by IVIS. **f,** Log-transformed *ex vivo* brain metastatic burden at endpoint, measured by IVIS and images of all brains. **g,** Quantification of metastases in brains via H&E. **h,** Representative H&E stains of metastases from two animals per group. **i,** UCD65 GFPLuc+ cells were injected intracardially into aged (>52w) NSG mice and mice randomized to receive vehicle (1% CMC, n=9) or BGJ398 (30mg/kg) 5x/week PO starting at the time of injection (W0-14, n=9) or 10 weeks post-injection (W10-14, n=9). **j,** Log-transformed head metastatic burden measured by IVIS. **k,** Log-tranformed *ex vivo* brain metastatic burden at endpoint, measured by IVIS. **l,** Quantification of metastases in brains via H&E. **m,** schematic of proposed pathway. For **e,j,** data were analyzed with 2-way ANOVA or mixed-effects analysis followed by Fisher’s LSD test. Significant p-values shown for last time point only. Gray dotted line denotes baseline IVIS signal from a tumor-free mouse. For **f,k,l,** data were analyzed with 1-way ANOVA followed by Fisher’s LSD test. Data in **g** were analyzed by Kruskal-Wallis and uncorrected Dunn’s tests.

Together, our data suggests a model whereby FGF2 and NCAM1-dependent FGFR1 activation of disseminated FGFR1-amplified ER+ BC cells drives neural/astrocytic interaction, migration and proliferation, resulting in FGFR1-dependent brain colonization. In young hosts, FGFR1 function is mainly driven by astrocytic FGF2, while in aged and E2-depleted mice, neuronal and astrocytic NCAM1 serve as non-canonical activators to drive FGFR1 function and ER+ BMs. Consistently, FGFR inhibitors can block early activation and prevent colonization, but not late-stage BCBM which are no longer dependent on this signaling (**Fig. 8m**).

## DISCUSSION

FGFR1-amplification has been shown to promote endocrine resistance in ER+ primary BC by maintaining ligand-independent ER transcription^16,17^, yet how the TME influences endocrine resistance at metastatic sites is less understood. Here we demonstrate that the increased BM potential of FGFR1+ ER+ BC cells results from interactions with canonical and non-canonical FGFR1 activation through astrocytes and neurons during early stages of BM colonization. Moreover, our data demonstrates that TME-mediated activation of FGFR1 in ER+ BCBM is critical for early but not late stages of BM colonization, which has profound implications for the use of FGFR inhibitors clinically. Specifically, our studies provide preclinical rationale for future clinical studies testing FGFR inhibitors in the prevention of BCBM.

Our results are in alignment with prior work showing that microenvironment-dependent activation of FGFR1 drives FGFR1-mediated ER signaling and endocrine therapy resistance in primary tumors^14^. FGFR ligands secreted by adipose tissue support ER+ primary tumor growth in obese post-menopausal mouse models^20^, and cells within the bone TME secrete FGF2 to promote ER+ BC cell plasticity^50^. However, our studies further demonstrate that the levels of FGFR1-activating ligands in the TME are key determinants of the dependency of ER+ BC cells for FGFR1 activation and response to FGFR inhibition. Furthermore, the contact-dependent nature of brain-mediated FGFR1 activation suggests that receptor inhibition or disruption of cell-cell interactions are the most relevant clinical targets for early brain colonization of ER+ BC. While the tumorigenic effect of FGFR1 amplification has been attributed to co-amplified oncogenes in the 8p region^51,52^, our FGFR1 KD studies demonstrate its specific role for brain colonization in aged and young hosts. The fact that FGFR1 KD differentially impacted extracranial metastases in aged mice, but not in young/E2 treated mice, further highlights the importance of studying ER+ BC in aged/E2 depleted mice and the context-dependent mechanisms influencing the progression of ER+ metastases.

Multiple studies have shown that interactions between disseminated cancer cells and astrocytes can eliminate or support the establishment of brain metastases^53–58^. While these studies have been performed with rapidly growing and aggressive cancer subtypes, our studies are the first to demonstrate brain-specific regulation of ER+ BMs in models that mimic the slow progression of their metastases. Our data showed that young and old astrocytes have different abilities to promote growth of ER+ BC cells, in part through differential expression of FGF2. However, the fact that conditioned media from astrocytes (lacking soluble FGF2) can activate pro-survival signaling in cancer cells in an FGFR1-independent manner, suggests that additional ligands secreted from astrocytes and other cells in the TME are likely to contribute to the progression of ER+ BMs, even those lacking FGFR1 amplification. It is possible that these signals from the brain TME contribute to the observed FGFR1-independence of late-stage brain metastases. Further studies are needed to define how additional astrocytic-mediated support of ER BM+ can contribute to late-stage ER+ BCBM progression.

Consistent with findings in rats^59,60^, our studies show for the first time that E2-suppression and aging downregulate FGF2 levels in the brain niche and directly impact FGFR1 activation. While these differences were initially observed in animals with late-stage BCBM, tumor-naïve mice showed similar FGF2 decreases with age. We showed the novel finding that NCAM1/FGFR1 interactions lead to FGFR1 kinase signaling activation and promote proliferation, invasive and migratory phenotypes in ER+ BC cells. While FGF2 decreases with aging and E2-depletion, NCAM1 expression remained unchanged, suggesting that FGFR1/NCAM interactions contribute to FGFR1 function throughout the progression of ER+ BCBM. Our findings that NCAM1 blockage decrease proliferation of cancer cells in brain slices from old, but not young mice, demonstrate a novel, context dependent mechanism controlling FGFR1 through the lifespan. Since the affinity of FGF for FGFR is approximately 10^6^ times higher than that of NCAM1 for FGFR^31,32^, higher FGF2 levels drive FGFR1 activation and function in young females. By contrast, NCAM1, which is constantly present cell surfaces at a much higher concentration (micromolar) than FGFs, emerged as an alternative driver in the aged/E2-depleted brain TME^30–32,43,61^. We recognize that our studies cannot fully address the individual contribution of brain FGF2 versus NCAM1 to FGFR1 function in young versus aged females, yet our findings align with studies in HeLa cells showing that despite similar kinase signaling activation, FGF2 and NCAM1 can result in different cellular responses^44,62^. The extent to which FGF2 and NCAM induce differential levels of pFGFR1 or trigger different recycling of FGFR from the cell membrane may explain the slower progression of metastases in older mice and the differential susceptibility of young and aged hosts to kinase inhibitors^44^.

Seminal studies demonstrated that brain metastatic growth of cancer cells depends on their ability to form synapses with neurons^47,48^, similar to findings in glioma cells^63,64^. Additional studies suggest that formation of synaptic structures involving presynaptic neurons, astrocytes, and tumor cells is a critical adaptation for BM colonization^46^. Our data showing that FGFR1 promotes increased synaptic puncta with neurons is suggestive of cancer-neuron synapse formation. While definitive proof that these interactions result in synaptic communication in ER+ BC cells is warranted, the formation of synaptic puncta and the migration of BC cells along neuronal tracks suggest that these cancer-neuron and FGFR1/NCAM1 interactions are implicated in the progression of ER+ BCBM. FGFR1 KD impairing ER+ BC cell migration along neurites further supports the notion that NCAM1/FGFR1 interaction is critical for the ability of ER+ BC cells to adhere to neurons and astrocytes, particularly during the early stages of BM colonization. FGFR1 KD UCD65 and UCD12 cells showed decreased brain signal (but not extracranial signal) immediately following intracardiac injection, suggesting that FGFR1 interactions with the brain TME are essential for early colonization or survival of disseminated cells upon arrival to the brain. Consistently, FGFR1 kinase inhibition decreased FGFR1 activation in early disseminated cancer cells and resulted in decreased number of micro metastases in both young and aged hosts, highlighting the potential value for FGFR1 inhibition in the prevention of BMs.

Our studies also show that late-stage FGFR1+ ER+ BMs are no longer dependent on FGFR1 activation. If this FGFR1-independence in large tumors occurs also in primary tumors or other metastatic sites, there could be an explanation for why FGFR1 amplification does not predict response to this therapy^65^. Further studies are needed to determine whether other FGFR1 inhibitors or increased brain-penetrance of these drugs may be more effective in a therapeutic setting^66^. Collectively, our work demonstrates that changes in the brain TME mediate FGFR1-amplified ER+ BCBM progression through interactions with astrocytes and neurons, which are critical for early brain metastatic colonization but not late progression. Importantly, our studies caution that current strategies to block FGFR1 function are ineffective to block non-canonical functions of FGFR1, particularly in elderly/E2-suppressed patients, and suggest that accounting for host-specific FGFR1 activation may better predict response to FGFR inhibitors than amplification status alone.

## MATERIALS AND METHODS

### Ethics Statement

All animal experiments and procedures were performed in an AAALAC-accredited facility and approved by the University of Colorado Denver | Anschutz Medical Campus Institutional Animal Care and Use Committee and by the DOD Animal Care and Use Review Office. Archival de-identified brain metastasis human samples were obtained under approved IRB protocols at the University of Colorado.

### Patient samples

ER+ BCBM (n = 45), including ER⁺/HER2⁻ (n = 30), ER⁺/HER2⁺ (n = 14), and ER⁺/PR⁺/HER2⁺ (n = 1) from archival de-identified human brain metastases samples were obtained under secondary use Colorado Multiple Institutional Review Board (COMIRB) protocol 23-1713 from the University of Colorado tissue biorepository (waiver of consent COMIRB protocol 15-1461).

### Cell Lines and Culture

ER^Low^-TN UCD46 and ER+ UCD4, UCD12, and UCD65 cell lines^38^ were cultured in DMEM/F12 with 10% FBS, P/S, 100ng/mL cholera toxin, and 1nM insulin. MCF7 (FGFR1-OE and EV) were cultured in MEM with EBSS, 10% FBS, P/S, 1x MEM amino acid solution, 1x MEM non-essential amino acids, and 1mM sodium pyruvate. SH-SY5Y cells were purchased from the Cell Technologies Shared Resource (RRID: SCR_021982) and maintained in DMEM/F12 with 10% FBS and P/S. Human astrocytes (Cat. #1800) were purchased from ScienceCell^TM^ Research Laboratories and maintained in astrocyte medium (AM, Cat. #1801) in flasks coated with Poly-L-Lysine (1ug/ml). Estrogen-free variations of all medias were created using phenol red-free media base and 2-5% charcoal-stripped FBS. Starvation media were created using phenol red-free media, no FBS, and 0.1% BSA.

### *In vivo* modeling of ER+ BC metastases

All *in vivo* studies were completed in NOD-*scid* IL2Rg^null^ (NSG) mice. Since circulating levels of E2 in gonadal intact young-female mice (2.7± 1 pg/ml) are similar to those of post-menopausal women (1.3-4.9 pg/ml)^10^, we mimicked post-menopausal BC using naïve (non-operated) or sham-operated aged females^11^. To study brain metastatic colonization, we use well recognized experimental models of metastasis, where cells are injected intracardially or in the carotid artery to bypass the exit from primary tumor and allow studies of seeding and metastatic colonization in the brain^67–69^. 500,000 cells were injected into the left ventricle of female NSG mice. Intracarotid artery injections were performed as previously described^67^. Spontaneous metastasis models were generated by injecting cells into the mammary fat pad of OVX NSG mice supplemented with E2. Tumors were removed after reaching 1 cm^3^, and mice were randomized into indicated groups. Injection success and luciferase-transfected cell growth were tracked via live bioluminescence imaging (BLI) using an IVIS® Spectrum optical scanner following subcutaneous injection of luciferin substrate (D-Luciferin, Potassium Salt, GoldBio 115144-35-9). For spontaneous models, the primary tumor site was covered during BLI measurements. BLI measurements included tumor-free mice to establish background. For mice treated with BGJ398 (Selleck Chemicals S2183), aliquots of drug were prepared daily and delivered via oral gavage of 100µL in 1% CMC. Drug and vehicle were provided 5 times weekly.

### Histological Quantification of Brain Metastases

Intraparenchymal BM were quantified as previously described^57^. Briefly, six hematoxylin and eosin (H&E)–stained sagittal brain sections (20μm thick) were examined per mouse. Sections were collected from the right hemisphere at 300μm intervals and analyzed using an ocular grid with a 4× objective. Micrometastases (defined as lesions measuring ≤300μm along their longest axis), and macrometastases (>300μm) were counted in each section by an investigator blinded to treatment group. Metastatic clusters were defined as groups of one or more micro or macrometastases in close spatial proximity as a readout of likely independent metastatic lesions. Numbers shown are the sum of counts from the six sections.

### Magnetic Resonance Imaging (MRI) and Analysis

To assess metastatic spread non-invasively, mice underwent MRI 16 weeks after tumor cell injection. Brain MRI was performed on anesthetized mice (1.5% isoflurane) using a 9.4 Tesla BioSpec animal MRI scanner (Bruker Medical, Billerica, MA) equipped with a 4-chanel phase-array mouse brain coil at the Colorado Animal Imaging Shared Resource (AISR). Each MRI session consisted of a tri-pilot localizer, followed by a high resolution T2-weighted fast spin echo (T2-turboRARE: field of view = 3 × 3 cm; repetition time = 4000ms; echo time = 80ms; slice thickness = 0.7 mm; 256 × 256 matrix size; 4 averages) with a 52-micron in-plane resolution^70,71^.

Total acquisition time for all sequences was 12 min 21 sec. The total metastatic gross volumes were quantified by placing hand-driven region of interest (ROI) on sequential axial and/ or sagittal T2-turboRARE images, summing total ROI areas and multiplying by 0.7 mm slice thickness (reported in mm3). The largest diameters of larger metastatic lesions were also reported for axial MR images. All MR image acquisition and analysis were performed using Bruker ParaVision 360neo v3.3 software. All image analysis was performed by an MRI physicist with >15 years of experience (NJS).

### Immunoblots

Membranes were blocked with 3% BSA in TBST and probed overnight with antibodies against FGFR1 (Abcam ab76464), pFGFR1 (Y654, Abcam ab59194; Y653/Y654, Cell Signaling 52928S), Akt (Cell Signaling 9272S), pAkt (S473, Cell Signaling 4060S), ERK (Cell Signaling 9102S), pERK (T202/T204, Cell Signaling 9101S), and NCAM1 (R&D AF2408), with GAPDH (Cell Signaling 97166S) or Tubulin (Sigma T5168) as loading controls. Membranes were washed with TBST and probed with secondary antibodies (Invitrogen A11371, A21109, A21058, A21084, A11374) for 1-2hrs. Blots were imaged on LiCor Odessey CLx and analyzed with ImageStudio Version 5.0 and 6.0. For multiple stains, blots were stripped (Millipore 2504) and reprobed as above.

### qRT-PCR

RNA was isolated with Trizol Reagent (Invitrogen 15596026) and reverse transcribed with Verso cDNA Synthesis Kit (Thermo AB-1453). cDNA was used to perform qPCR with PowerUp SYBR Green Master Mix (Applied Biosystems A25742) on Applied Biosystems QuantStudio 6 Flex. Gene expression was calculated via ddCt. Primer Sequences:

*FGFR1* (F: AACCTGACCACAGAATTGGAGGCT, R: TGCTGCCGTACTCATTCTCCACA)

*NCAM1* (F: AATTTACCGCGGCAAGAACATC, R: CCTGGCTGGGAACAATATCCAC)

*RPLP0* (F: GTGATGTGCAGCTGATCAAGACT, R: GATGACCAGCCCAAAGGAGA)

### Primary Astrocyte Culture and Co-culture

Brains from P0-P1 CD1 mouse pups were dissected, and primary astrocytes were isolated and cultured as previously described^72^. For co-culture experiments, 50,000 or 10,000 cancer cells were seeded into 6-well or 24-well plates coated with laminin (1µg/ml), respectively, onto astrocyte monolayers at 100% confluence in E2-free medium (DMEM/F12 phenol red-free, supplemented with 5% charcoal-stripped FBS, 0.1% BSA, 1x penicillin/streptomycin, 1x non- essential amino acids, 1x sodium pyruvate, and 1 nM insulin). Aged mouse astrocytes (aged-mAst) were generated by maintaining cultures in growth medium (DMEM + 10% FBS, 1% P/S) for three months prior to co-culture^73^. In contrast, mAst P0 refers to astrocytes maintained *in vitro* for only 8 days. For co-culture with human astrocytes, cells were seeded under the same conditions as murine astrocytes. To generate 10x or 20x concentrated astrocyte-conditioned media (CM), confluent astrocyte cultures were incubated for 72 hours in serum-free, phenol red-free DMEM/F12. Supernatants were then collected and concentrated using 3,000Da molecular weight centrifugal filters.

### Isolation of Primary Cells from Adult Mouse Brain

Whole brain tissue was dissected from young (12 weeks) and aged (>56 weeks) female NSG mice following PBS cardiac perfusion. Cell suspensions were obtained via enzymatic dissociation using the Papain Dissociation System (Worthington Biochemical, LK003150), followed by magnetic-activated cell sorting (MACS; Miltenyi Biotec) according to the manufacturer’s instructions. Cell-type-specific isolation was performed using the following reagents from Miltenyi Biotec: Myelin Removal Beads II (130-096-733) for oligodendrocytes, CD11b MicroBeads (130-093-634) for microglia, Anti-ACSA-2 MicroBeads (130-097-678) for astrocytes, and the Neuron Isolation Kit (130-115-389) for neurons.

### FGF2 ELISA

Levels of FGF2 were quantified using the Mouse/Rat FGF basic/FGF2/bFGF Quantikine ELISA Kit (R&D Systems, MFB00) according to the manufacturer’s instructions.

### Lentivirus and AAV

The following shRNA plasmids were purchased from the Functional Genomics Shared Resources (RRID:SCR_021987): TRCN0000121186, TRCN0000000419, and TRCN0000000418 targeting human FGFR1; TRCN0000373085 and TRCN0000373034 targeting NCAM1, TRCN0000355839 targeting Fgf2. MISSION® shRNA Plasmid DNA Control Vectors SHC216 and SCH002 were purchased from Sigma. Lentiviral particles were produced in 293-T cells and concentrated via ultracentrifugation. Viral particles were added to all cells with 6µg/mL polybrene and cancer cells were selected with 2µg/mL puromycin. For transduction of astrocytes, lentivirus was not concentrated and cells were not selected. AAV shRNA particles were purchased from PackGene and designed to include CMV-mCherry as a reporter and the same shRNA sequences as TRCN0000355839 (Fgf2) and TRCN0000373085 (Ncam1). Control AAV was purchased from AAVnerGene (DP001006). Astrocytes were infected with 10,000 viral particles per cell.

### Organotypic Co-culture

GFP+ cancer cells were cultured in ultra-low attachment plates to form spheroids. NSG mice were sacrificed and brains excised, sectioned into 250µm coronal slices, and collected in 1x HBSS with 2.5mM HEPES, 30mM D-Glucose, 1mM CaCl2, 1mM MgSO4, and 4mM NaHCO3 (Complete HBSS). Slices were plated in 12-well transwells. 1mL of media containing 25% complete HBSS, 25% donor equine serum, 50% MEM, 1x GlutaMAX (Gibco 35050061), and P/S was added to the well below the transwell. Spheroids of cancer cells were plated on top of the brain slices in <2µL media. Spheres were allowed to settle for 24 hours and images were acquired via stereo fluorescent microscopy every 48 hours. Growth was quantified via GFP intensity corrected for background autofluorescence of adjacent brain tissue (ImageJ). For treatments of organotypic brain slices with IgG (Sigma M5284), αFGF2 (Antibody System RMC38201), or αNCAM1(Sigma AB5032) neutralizing antibodies, 0.5µg was added on top of each slice in 10µL, with an additional 0.5µg added to the media in the well below. Spheroids of UCD12 cells were plated 24 hours after antibody treatment.

### Immunohistochemistry and Immunofluorescence

Formalin-fixed, paraffin-embedded sections of ER⁺ BCBM from patient samples obtained at the University of Colorado were deparaffinized in xylene and rehydrated through a graded ethanol series to distilled water. Heat-induced antigen retrieval was performed in 10 mM citrate buffer (pH 6.0) or Tris-EDTA buffer (pH 9.0) at 125 °C and 25 psi for 10 minutes. Endogenous peroxidase activity was quenched with 3% hydrogen peroxide, and sections were blocked with 10% normal horse serum (ImmPRESS, Vector Laboratories). Slides were incubated overnight at RT with primary antibodies against FGFR1 (Invitrogen PA5-25979) or pFGFR1 (Y654, Abcam ab59194), followed by HRP-conjugated anti-rabbit IgG (ImmPRESS, Vector Laboratories). Signal was developed using 3,3′-diaminobenzidine (DAB; DAKO). Slides were scanned using an Aperio ScanScope T3 (Leica Biosystems), and membrane expression of FGFR1 and pFGFR1 in tumor cells was quantified using the Aperio Image Analysis software.

For immunofluorescence, cells grown on coverslips and free-floating brain slices were fixed with 4% paraformaldehyde (PFA) and heat-induced antigen retrieval was performed in boiling 10 mM citrate buffer (pH 6.0). Coverslips, slices, or deparaffinized formalin-fixed, paraffin-embedded sections (as described above), were blocked with 10% normal donkey serum (NDS) in TBST for 1 hour at RT. Primary antibodies against GFAP (Invitrogen 12-0300), FGF2 (Novusbio NBP3-15367), pFGFR1 (Y654, Abcam ab59194), NCAM1 (R&D AF2408), PanCK (DAKO M0821), Synapsin1/2 (Synaptic Systems 106004), and PSD95 (Invitrogen MA1-045) were incubated for 2 hours or overnight at RT. For double staining, primary antibodies were applied either sequentially in separate incubations or pooled. After washing with PBS containing 0.01% Triton X-100, coverslips, slices, or slides were incubated with secondary antibodies (Jackson ImmunoResearch 712-545-153, 712-585-153, 706-165-148, 715-545- 150, 715-585-150, 715-605-150, 705-585-147, 711-545-152, 711-585-152, 711-605-152) diluted in TBST for 2 hours at RT in the dark. Nuclei were stained with DAPI (1 µg/mL in methanol or PBS) for 10 minutes at RT and samples were mounted with FluoroMount-G (Southern Biotech).

### Growth Assays

Cells were maintained in E2-free media for 48 hours before being trypsinized with phenol red-free trypsin and plated at densities of 2,000 MCF7, 10,000 UCD12, and 20,000 UCD65 cells per well of duplicate 96-well plates in 100µL of media. Following 24 hours, 100µL of media containing each treatment (2x) was added to the wells of one plate. The duplicate plate was fixed with 10% TCA. Growth was tracked via Incucyte S3 system where 10x phase contrast and GFP images were obtained every 12 hours. Treatments were maintained by regularly changing 50% of media. At the conclusion of the Incucyte tracking, plates were fixed with 10% TCA. Fixed plates were stained with 0.04% SRB (Sigma) and total protein was quantified via plate reader measurements at 565, 525, and 490nm subtracting 690nm background. Proliferation was additionally measured by quantifying fold change of SRB signal between the two plates.

### Digital Spatial Transcriptomics

Female NSG mice were ovariectomized or sham operated and 15 days later injected via intracarotid artery with 250,000 UCD65 GFPLuc+ cells. Mice were classified according to their age as young (between 7 and 15 weeks) or aged (>60 weeks). Development of brain metastasis was monitored during 15 weeks via IVIS. Six months after injection, animals were euthanized by ketamine/xylazine overdose and perfused with PBS/4%PFA. Brains were cryopreserved in 20% sucrose and stored at 4°C. Paraffin embedded fixed coronal sections were stained with antibodies targeting PanCK (DAKO M0821), Iba1 (WAKO PTG5394), GFAP (Invitrogen 13-0300) and DAPI. One set of brain sections from young-sham, aged-sham, young-OVX and aged-OVX mice (n = 1 animal per condition) were probed by in situ hybridization with the GeoMx® Mouse Whole Transcriptome Atlas (WTA) library. Then, using the GeoMX Digital Spatial Profiler (DSP), barcoded RNA-probes in the Iba+/GFAP+ regions of interest (ROI) within a maximum of 400µm from PanCK+ BCBM were UV-cleaved and processed by Illumina sequencing. Similarly, another set of brain sections from young-sham and aged-sham mice were probed by in situ hybridization with the GeoMx® Human WTA library. Then, with the GeoMX DSP, RNA-probes in the PanCK+ ROIs (human breast cancer cells) were UV-cleaved and processed by Illumina sequencing. A total of 48 Iba+/GFAP+ ROIs (young-sham = 18, aged-sham = 16, young-OVX = 9, and aged-OVX = 5) and 59 PanCK+ ROIs (aged-sham = 16 and young-sham = 43) were retrieved and sequenced using an Illumina NovaSeq X plus sequencer in one lane on a 10B flow cell at 2×150bp configuration. Sequencing was performed by the Genomics Shared Resource (RRID: SCR_021984).

Using the GeoMx® NGS Pipeline (Nanostring) and the Probe Kit Configuration (PKC) files for Human or Mouse RNA WTA, the FASTQ libraries were processed to obtain the Digital Count Conversion (DCC) files. DCC files were analyzed in R (4.4.2) with NanoStringNCTools^74^, GeomxTools^75^ and GeoMxWorkflows^76^ packages. All ROIs with gene detection rate (defined as the number of genes in the sample divided by the number of genes in the WTA) above 1%, more than 100 nuclei, and all probes in the highest quartile above the background reading were retained for further analysis (**Supplementary Fig. 3**). Raw counts were normalized using the Q3 method in NanoStringNCTools. Normalized counts were used to UMAP analysis with the umap package in R. Differential expression analyses were performed with limma-voom in standR^77,78^ using the GeoMXAnalysisWorkflow package in R^79^. Differentially expressed genes were visualized as MA plots with ggplot2.

Log-CPM (counts per million) values were transformed with “voom” to get the appropriate observation-level weights and fitted to a linear model with “fit”. Then using “contrasts.fit”, the array was used to get estimated coefficients and standard errors for each pairwise comparison. Finally, the differential expression statistics were computed using empirical Bayes moderation with “ebayes”. Differentially expressed genes lists were used to perform a gene sets enrichment analysis (GSEA) following the GeoMXAnalysisWorkflow method^79^. Briefly, MSigDB Hallmarks, MSigDB C2, Gene Ontology and KEGG databases were downloaded in R using the msigdb and GSEABase libraries^80,81^. Enrichment analyses were performed using fry from the limma package^78^. Finally, using vissE package^82^ we clustered significant (FDR <0.05) enriched gene sets based on the genes overlap with a threshold of 35%. Clusters were further analyzed by text mining to identify common biological patterns using the gene set names. Additionally, frequency of each gene in each cluster was plotted vs their respective log2FC. Clusters were ranked by size and FDR and top 9 clusters were plotted using igraph^83^.

### Recombinant protein treatment for downstream signaling

1 million UCD65, UCD12, or MCF7 cells were plated in 60mm plates. 12 hours after plating, cells were washed three times with PBS and media was replaced with starvation media. Following 48-72 hours in starvation media, media was replaced with starvation media containing the indicated concentrations of FGF2 (PeproTech AF-100-18B) or NCAM1 (Acro Biosystems NC1-H5223). Pretreatments with BGJ398 (Selleck Chemicals S2183) were completed 1 hour prior to treatments. Following treatment, protein was collected in RIPA buffer.

### RNA Sequencing

UCD65 and UCD12 cells were serum-starved and then treated with vehicle, 10ng/mL FGF2, or 20ug/mL for 24hr. RNA was isolated with Trizol Reagent (Invitrogen 15596026) followed by RNeasy Cleanup Kit (Qiagen 74204). mRNA libraries were created using poly-A enrichment and sequenced at 40M PE150 reads by NovaSeq X Plus Series (Illumina). Reads were filtered by quality (removal of reads containing adaptors, >10% undetermined bases, and QScore <5 for 50% of bases) and aligned with HiSAT2 (2.2.1) to Homo Sapiens Reference Genome (GRCh38/hg38). Raw counts were obtained with featureCounts (2.0.6), and analyzed in R 4.5.0. Normalized counts and differential expression were obtained with deseq2^84^ and used to perform gene sets enrichment analysis (GSEA) as described above for digital spatial transcriptomics, clustered based on 25% gene overlap.

### Primary Neuron Culture and Co-Culture

Hippocampi were dissected from P0-1 CD1 pups and neurons were isolated as previously described^85^. Primary neurons were cultured on poly-l-lysine coated coverslips in 24-well plates and maintained in phenol red-free neurobasal media supplemented with GlutaMAX, 1x B27, and P/S, with 50% media replaced twice weekly. Five days after isolation, 1.5µM cytosine arabinoside was added to each well. After 48 hours, 50% media was replaced. For co-cultures, 5,000-10,000 cancer cells were added to each well 24 hours after cytosine arabinoside removal. Co-cultures grew for 5-10 additional days before coverslips were fixed.

### Synapse Detection and Quantification

Synapsin1/2 and PSD95 were costained via immunofluorescence and samples were imaged via confocal microscopy (3I MARIANAS Inverted Spinning Disk) at 40x magnification. SynQuant ImageJ plugin^86^ was used to quantify synapses as punctate co-localization of the two markers, and synapse number was controlled for analysis area differences between images.

### Neural Migration Analysis

Phase and GFP images were collected every 12 hours. GFP+ ROIs were defined to indicate cells in each image. Using ImageJ, phase images were used to stabilize motion across time points^87^, and these corrections were applied to the GFP+ ROIs. Movement was tracked using TrackMate^88,89^.

### Statistical Analysis and Reproducibility

Statistical analyses are specified in figure legends and were performed with GraphPad Prism 10.4.2 for predetermined comparisons only. All tests were 2-tailed. Statistical analysis of sequencing data was performed in R 4.4.2 and 4.5.0. For animal experiments involving injections of different cells, groups were littermate-matched and injections were blinded. For treatment of established tumors, groups were randomized. In studies involving aged mice, there was expected age-related, non-tumor related mortality. These animals were excluded from final analysis and initial group sizes were 20% larger to account for this loss. Where possible, analyses were blinded to experimental conditions or performed using automated functions to reduce bias.

## Data Availability

Digital Spatial Profiling and RNA-sequencing data will be uploaded to NCBI’s Gene Expression Omnibus prior to publication.

## Code Availability

All information is provided in the Methods, including packages, software, and pipelines. No custom code was generated during these studies. Source Data are provided with this paper.

## Acknowledgements/Grant Support

This study was supported by DoD-BCRP W81XWH-22-1-0042 (DMC), R37CA227984 (DMC), T32CA190216 (MSF), F31CA294726 (MSF), University of Colorado Cancer Center Pilot Funds (DMC, CAS), the University of Colorado Cancer Center Support Grant (P30CA046934), and NIH High-End and SIFAR S10 Shared Instrumentation Grants (S10OD023485 and S10OD027023). PK is supported by R01CA258766. We would like to thank the Biorepository Core Facility and personnel at the CU Brain Tumor Biorepository for providing de-identified human tissues, Animal Imaging Shared Resource and Jenna Steiner, Genomics Shared Resource and Schuyler Lee, Functional Genomics Shared Resource, Human Immune Monitoring Shared Resource and Kimberly Jordan, and the Cell Technologies Shared Resource for their support of these studies. Imaging was performed in the Advanced Light Microscopy Core facility of the NeuroTechnology Center at the University of Colorado Anschutz Medical Campus, which is supported in part by Diabetes Research Center Grant P30DK116073. We would also like to thank Lianne Kraemer, Julia Maués, and Christine Hodgdon for their input and support.

## Authorship contribution

Study supervision: DMC; Conception and design: DMC, MSF; Data acquisition, analysis, resources: DMC, MSF, JAJ, RAM, KLFA, MJC, TCP, ENB, SNK, PK, NJS, CAS, EAW. All authors contributed to writing and approved manuscript.

## REFERENCES

1 Kaal, E. C. A., Niël, C. G. J. H. & Vecht, C. J. Therapeutic management of brain metastasis. The Lancet Neurology 4, 289–298 (2005). 10.1016/S1474-4422(05)70072-7

2 Palmieri, D. et al. Brain metastases of breast cancer. Breast disease 26, 139–147 (2006).

3 Morris, P. G. et al. Limited Overall Survival in Patients with Brain Metastases from Triple Negative Breast Cancer. The Breast Journal 18, 345–350 (2012). 10.1111/j.1524-4741.2012.01246.x

4 Press, D. J., Miller, M. E., Liederbach, E., Yao, K. & Huo, D. De novo metastasis in breast cancer: occurrence and overall survival stratified by molecular subtype. Clin Exp Metastasis 34, 457–465 (2017). 10.1007/s10585-017-9871-9

5 Kennecke, H. et al. Metastatic behavior of breast cancer subtypes. J Clin Oncol 28, 3271–3277 (2010). 10.1200/JCO.2009.25.9820

6 Heitz, F. et al. Triple-negative and HER2-overexpressing breast cancers exhibit an elevated risk and an earlier occurrence of cerebral metastases. Eur J Cancer 45, 2792–2798 (2009). 10.1016/j.ejca.2009.06.027

7 Chatzistamou, I. & Kiaris, H. Modeling estrogen receptor-positive breast cancers in mice: is it the best we can do? Endocr Relat Cancer 23, C9–C12 (2016). 10.1530/ERC-16-0397

8 Sledge, G. W. et al. Past, present, and future challenges in breast cancer treatment. J Clin Oncol 32, 1979–1986 (2014). 10.1200/JCO.2014.55.4139

9 Turrell, F. K. et al. Age-associated microenvironmental changes highlight the role of PDGF-C in ER(+) breast cancer metastatic relapse. Nat Cancer 4, 468–484 (2023). 10.1038/s43018-023-00525-y

10 Rothman, M. S. et al. Reexamination of testosterone, dihydrotestosterone, estradiol and estrone levels across the menstrual cycle and in postmenopausal women measured by liquid chromatography-tandem mass spectrometry. Steroids 76, 177–182 (2011). 10.1016/j.steroids.2010.10.010

11 Osborne, C. K., Hobbs, K. & Clark, G. M. Effect of estrogens and antiestrogens on growth of human breast cancer cells in athymic nude mice. Cancer Res 45, 584–590 (1985).

12 Offner, H. & Polanczyk, M. A potential role for estrogen in experimental autoimmune encephalomyelitis and multiple sclerosis. Ann N Y Acad Sci 1089, 343–372 (2006). 10.1196/annals.1386.021

13 Guerini-Rocco, E. et al. Genomic Aberrations and Late Recurrence in Postmenopausal Women with Hormone Receptor-positive Early Breast Cancer: Results from the SOLE Trial. Clin Cancer Res (2021). 10.1158/1078-0432.CCR-20-0126

14 DiGiacomo, J. W., Godet, I., Trautmann-Rodriguez, M. & Gilkes, D. M. Extracellular Matrix-Bound FGF2 Mediates Estrogen Receptor Signaling and Therapeutic Response in Breast Cancer. Mol Cancer Res 19, 136–149 (2021). 10.1158/1541-7786.MCR-20-0554

15 Garcia-Recio, S. et al. FGFR4 regulates tumor subtype differentiation in luminal breast cancer and metastatic disease. J Clin Invest 130, 4871–4887 (2020). 10.1172/JCI130323

16 Formisano, L. et al. Aberrant FGFR signaling mediates resistance to CDK4/6 inhibitors in ER+ breast cancer. Nat Commun 10, 1373 (2019). 10.1038/s41467-019-09068-2

17 Drago, J. Z. et al. FGFR1 Amplification Mediates Endocrine Resistance but Retains TORC Sensitivity in Metastatic Hormone Receptor-Positive (HR(+)) Breast Cancer. Clin Cancer Res 25, 6443–6451 (2019). 10.1158/1078-0432.CCR-19-0138

18 Ding, K. et al. FGFR4 in endocrine resistance: overexpression and estrogen regulation without direct causative role. Breast Cancer Res Treat 211, 501–515 (2025). 10.1007/s10549-025-07666-x

19 Xie, N. et al. FGFR aberrations increase the risk of brain metastases and predict poor prognosis in metastatic breast cancer patients. Ther Adv Med Oncol 12, 1758835920915305 (2020). 10.1177/1758835920915305

20 Wellberg, E. A. et al. FGFR1 underlies obesity-associated progression of estrogen receptor-positive breast cancer after estrogen deprivation. JCI Insight 3 (2018). 10.1172/jci.insight.120594

21 Sytnyk, V., Leshchyns’ka, I. & Schachner, M. Neural Cell Adhesion Molecules of the Immunoglobulin Superfamily Regulate Synapse Formation, Maintenance, and Function. Trends Neurosci 40, 295–308 (2017). 10.1016/j.tins.2017.03.003

22 Dabrowski, A., Terauchi, A., Strong, C. & Umemori, H. Distinct sets of FGF receptors sculpt excitatory and inhibitory synaptogenesis. Development 142, 1818–1830 (2015). 10.1242/dev.115568

23 Terauchi, A. et al. Distinct FGFs promote differentiation of excitatory and inhibitory synapses. Nature 465, 783–787 (2010). 10.1038/nature09041

24 Umemori, H., Linhoff, M. W., Ornitz, D. M. & Sanes, J. R. FGF22 and its close relatives are presynaptic organizing molecules in the mammalian brain. Cell 118, 257–270 (2004). 10.1016/j.cell.2004.06.025

25 Klimaschewski, L. & Claus, P. Fibroblast Growth Factor Signalling in the Diseased Nervous System. Mol Neurobiol 58, 3884–3902 (2021). 10.1007/s12035-021-02367-0

26 Ohkubo, Y., Uchida, A. O., Shin, D., Partanen, J. & Vaccarino, F. M. Fibroblast growth factor receptor 1 is required for the proliferation of hippocampal progenitor cells and for hippocampal growth in mouse. J Neurosci 24, 6057–6069 (2004). 10.1523/JNEUROSCI.1140-04.2004

27 Gonzalez, A. M., Berry, M., Maher, P. A., Logan, A. & Baird, A. A comprehensive analysis of the distribution of FGF-2 and FGFR1 in the rat brain. Brain Res 701, 201–226 (1995). 10.1016/0006-8993(95)01002-x

28 Ma, D. K., Ponnusamy, K., Song, M. R., Ming, G. L. & Song, H. Molecular genetic analysis of FGFR1 signalling reveals distinct roles of MAPK and PLCgamma1 activation for self-renewal of adult neural stem cells. Mol Brain 2, 16 (2009). 10.1186/1756-6606-2-16

29 Cambon, K. et al. A synthetic neural cell adhesion molecule mimetic peptide promotes synaptogenesis, enhances presynaptic function, and facilitates memory consolidation. J Neurosci 24, 4197–4204 (2004). 10.1523/JNEUROSCI.0436-04.2004

30 Christensen, C., Berezin, V. & Bock, E. Neural cell adhesion molecule differentially interacts with isoforms of the fibroblast growth factor receptor. Neuroreport 22, 727–732 (2011). 10.1097/WNR.0b013e3283491682

31 Kiselyov, V. V., Soroka, V., Berezin, V. & Bock, E. Structural biology of NCAM homophilic binding and activation of FGFR. J Neurochem 94, 1169–1179 (2005). 10.1111/j.1471-4159.2005.03284.x

32 Kiselyov, V. V. et al. Structural basis for a direct interaction between FGFR1 and NCAM and evidence for a regulatory role of ATP. Structure 11, 691–701 (2003). 10.1016/s0969-2126(03)00096-0

33 Krushel, L. A., Tai, M. H., Cunningham, B. A., Edelman, G. M. & Crossin, K. L. Neural cell adhesion molecule (N-CAM) domains and intracellular signaling pathways involved in the inhibition of astrocyte proliferation. Proc Natl Acad Sci U S A 95, 2592–2596 (1998). 10.1073/pnas.95.5.2592

34 Kobayashi, S., Vidal, I., Pena, J. D. & Hernandez, M. R. Expression of neural cell adhesion molecule (NCAM) characterizes a subpopulation of type 1 astrocytes in human optic nerve head. Glia 20, 262–273 (1997). 10.1002/(sici)1098-1136(199707)20:3<262::aid-glia10>3.0.co;2-s

35 Weledji, E. P. & Assob, J. C. The ubiquitous neural cell adhesion molecule (N-CAM). Ann Med Surg (Lond) 3, 77–81 (2014). 10.1016/j.amsu.2014.06.014

36 Knafo, S. et al. Facilitation of AMPA receptor synaptic delivery as a molecular mechanism for cognitive enhancement. PLoS Biol 10, e1001262 (2012). 10.1371/journal.pbio.1001262

37 Leshchyns’ka, I., Sytnyk, V., Morrow, J. S. & Schachner, M. Neural cell adhesion molecule (NCAM) association with PKCbeta2 via betaI spectrin is implicated in NCAM-mediated neurite outgrowth. J Cell Biol 161, 625–639 (2003). 10.1083/jcb.200303020

38 Finlay-Schultz, J. et al. New generation breast cancer cell lines developed from patient-derived xenografts. Breast Cancer Res 22, 68 (2020). 10.1186/s13058-020-01300-y

39 Finlay-Schultz, J. et al. Breast Cancer Suppression by Progesterone Receptors Is Mediated by Their Modulation of Estrogen Receptors and RNA Polymerase III. Cancer Res 77, 4934–4946 (2017). 10.1158/0008-5472.CAN-16-3541

40 Beenken, A. & Mohammadi, M. The FGF family: biology, pathophysiology and therapy. Nat Rev Drug Discov 8, 235–253 (2009). 10.1038/nrd2792

41 Plotnikov, A. N., Schlessinger, J., Hubbard, S. R. & Mohammadi, M. Structural basis for FGF receptor dimerization and activation. Cell 98, 641–650 (1999). 10.1016/s0092-8674(00)80051-3

42 Schlessinger, J. et al. Crystal structure of a ternary FGF-FGFR-heparin complex reveals a dual role for heparin in FGFR binding and dimerization. Mol Cell 6, 743–750 (2000). 10.1016/s1097-2765(00)00073-3

43 Aonurm-Helm, A., Berezin, V., Bock, E. & Zharkovsky, A. NCAM-mimetic, FGL peptide, restores disrupted fibroblast growth factor receptor (FGFR) phosphorylation and FGFR mediated signaling in neural cell adhesion molecule (NCAM)-deficient mice. Brain Res 1309, 1–8 (2010). 10.1016/j.brainres.2009.11.003

44 Francavilla, C. et al. The binding of NCAM to FGFR1 induces a specific cellular response mediated by receptor trafficking. J Cell Biol 187, 1101–1116 (2009). 10.1083/jcb.200903030

45 Niethammer, P. et al. Cosignaling of NCAM via lipid rafts and the FGF receptor is required for neuritogenesis. J Cell Biol 157, 521–532 (2002). 10.1083/jcb.200109059

46 Zeng, Q. et al. Synaptic proximity enables NMDAR signalling to promote brain metastasis. Nature 573, 526–531 (2019). 10.1038/s41586-019-1576-6

47 Venkataramani, V. et al. Direct excitatory synapses between neurons and tumor cells drive brain metastatic seeding of breast cancer and melanoma. bioRxiv, 2024.2001.2008.574608 (2024). 10.1101/2024.01.08.574608

48 Savchuk, S. et al. Neuronal activity-dependent mechanisms of small cell lung cancer pathogenesis. Nature 646, 1232–1242 (2025). 10.1038/s41586-025-09492-z

49 Lassman, A. B. et al. Infigratinib in Patients with Recurrent Gliomas and FGFR Alterations: A Multicenter Phase II Study. Clin Cancer Res 28, 2270–2277 (2022). 10.1158/1078-0432.CCR-21-2664

50 Bado, I. L. et al. The bone microenvironment increases phenotypic plasticity of ER(+) breast cancer cells. Dev Cell 56, 1100–1117 e1109 (2021). 10.1016/j.devcel.2021.03.008

51 Ray, M. E. et al. Genomic and expression analysis of the 8p11-12 amplicon in human breast cancer cell lines. Cancer Res 64, 40–47 (2004). 10.1158/0008-5472.can-03-1022

52 Voutsadakis, I. A. 8p11.23 Amplification in Breast Cancer: Molecular Characteristics, Prognosis and Targeted Therapy. J Clin Med 9 (2020). 10.3390/jcm9103079

53 Chen, Q. et al. Carcinoma-astrocyte gap junctions promote brain metastasis by cGAMP transfer. Nature 533, 493–498 (2016). 10.1038/nature18268

54 Zhang, L. et al. Microenvironment-induced PTEN loss by exosomal microRNA primes brain metastasis outgrowth. Nature 527, 100–104 (2015). 10.1038/nature15376

55 Valiente, M. et al. Serpins promote cancer cell survival and vascular co-option in brain metastasis. Cell 156, 1002–1016 (2014). 10.1016/j.cell.2014.01.040

56 Kienast, Y. et al. Real-time imaging reveals the single steps of brain metastasis formation. Nat Med 16, 116–122 (2010). 10.1038/nm.2072

57 Sartorius, C. A. et al. Estrogen promotes the brain metastatic colonization of triple negative breast cancer cells via an astrocyte-mediated paracrine mechanism. Oncogene 35, 2881–2892 (2016). 10.1038/onc.2015.353

58 Contreras-Zarate, M. J. et al. Estradiol induces BDNF/TrkB signaling in triple-negative breast cancer to promote brain metastases. Oncogene 38, 4685–4699 (2019). 10.1038/s41388-019-0756-z

59 Chadashvili, T. & Peterson, D. A. Cytoarchitecture of fibroblast growth factor receptor 2 (FGFR-2) immunoreactivity in astrocytes of neurogenic and non-neurogenic regions of the young adult and aged rat brain. J Comp Neurol 498, 1–15 (2006). 10.1002/cne.21009

60 Huang, C., Yuan, P., Wu, J. & Huang, J. Estrogen regulates excitatory amino acid carrier 1 (EAAC1) expression through sphingosine kinase 1 (SphK1) transacting FGFR-mediated ERK signaling in rat C6 astroglial cells. Neuroscience 319, 9–22 (2016). 10.1016/j.neuroscience.2016.01.027

61 Jurgenson, M., Aonurm-Helm, A. & Zharkovsky, A. Partial reduction in neural cell adhesion molecule (NCAM) in heterozygous mice induces depression-related behaviour without cognitive impairment. Brain Res 1447, 106–118 (2012). 10.1016/j.brainres.2012.01.056

62 Francavilla, C. et al. Neural cell adhesion molecule regulates the cellular response to fibroblast growth factor. J Cell Sci 120, 4388–4394 (2007). 10.1242/jcs.010744

63 Venkatesh, H. S. et al. Electrical and synaptic integration of glioma into neural circuits. Nature 573, 539–545 (2019). 10.1038/s41586-019-1563-y

64 Venkataramani, V. et al. Glutamatergic synaptic input to glioma cells drives brain tumour progression. Nature 573, 532–538 (2019). 10.1038/s41586-019-1564-x

65 Coombes, R. C. et al. Results of the phase IIa RADICAL trial of the FGFR inhibitor AZD4547 in endocrine resistant breast cancer. Nat Commun 13, 3246 (2022). 10.1038/s41467-022-30666-0

66 Knight, W. et al. Abstract 6296: Preclinical evaluation of a panel of FGFR inhibitors for their normal brain and brain tumor distribution. Cancer Research 82, 6296–6296 (2022). 10.1158/1538-7445.Am2022-6296

67 Zhang, C., Lowery, F. J. & Yu, D. Intracarotid Cancer Cell Injection to Produce Mouse Models of Brain Metastasis. J Vis Exp (2017). 10.3791/55085

68 Zhang, C. & Yu, D. Microenvironment determinants of brain metastasis. Cell Biosci 1, 8 (2011). 10.1186/2045-3701-1-8

69 Miarka, L. & Valiente, M. Animal models of brain metastasis. Neurooncol Adv 3, v144–v156 (2021). 10.1093/noajnl/vdab115

70 Koza, L. A. et al. Immunocal(R) limits gliosis in mouse models of repetitive mild-moderate traumatic brain injury. Brain Res 1808, 148338 (2023). 10.1016/j.brainres.2023.148338

71 Pierce, A. M. et al. Establishment of patient-derived orthotopic xenograft model of 1q+ posterior fossa group A ependymoma. Neuro Oncol 21, 1540–1551 (2019). 10.1093/neuonc/noz116

72 de Vellis, J. & Cole, R. Preparation of mixed glial cultures from postnatal rat brain. Methods Mol Biol 814, 49–59 (2012). 10.1007/978-1-61779-452-0_4

73 Pertusa, M., Garcia-Matas, S., Rodriguez-Farre, E., Sanfeliu, C. & Cristofol, R. Astrocytes aged in vitro show a decreased neuroprotective capacity. J Neurochem 101, 794–805 (2007). 10.1111/j.1471-4159.2006.04369.x

74 NanoStringNCTools: NanoString nCounter Tools (2024).

75 GeomxTools: NanoString GeoMx Tools (2024).

76 GeoMxWorkflows: GeoMx Digital Spatial Profiler (DSP) data analysis workflows (2024).

77 Liu, N. et al. standR: spatial transcriptomic analysis for GeoMx DSP data. Nucleic Acids Res 52, e2 (2024). 10.1093/nar/gkad1026

78 Ritchie, M. E. et al. limma powers differential expression analyses for RNA-sequencing and microarray studies. Nucleic Acids Res 43, e47 (2015). 10.1093/nar/gkv007

79 GeoMXAnalysisWorkflow: GeoMXAnalysisWorkflow (2023).

80 msigdb: An ExperimentHub Package for the Molecular Signatures Database (MSigDB) (2024).

81 GSEABase: Gene set enrichment data structures and methods (2024).

82 vissE: Visualising Set Enrichment Analysis Results (2024).

83 igraph: Network Analysis and Visualization in R (2025).

84 Love, M. I., Huber, W. & Anders, S. Moderated estimation of fold change and dispersion for RNA-seq data with DESeq2. Genome Biol 15, 550 (2014). 10.1186/s13059-014-0550-8

85 Beaudoin, G. M., 3rd et al. Culturing pyramidal neurons from the early postnatal mouse hippocampus and cortex. Nat Protoc 7, 1741–1754 (2012). 10.1038/nprot.2012.099

86 Wang, Y. et al. SynQuant: an automatic tool to quantify synapses from microscopy images. Bioinformatics 36, 1599–1606 (2020). 10.1093/bioinformatics/btz760

87 “The image stabilizer plugin for ImageJ,” https://www.cs.cmu.edu/∼kangli/code/Image_Stabilizer.html (2008).

88 Tinevez, J. Y. et al. TrackMate: An open and extensible platform for single-particle tracking. Methods 115, 80–90 (2017). 10.1016/j.ymeth.2016.09.016

89 Ershov, D. et al. TrackMate 7: integrating state-of-the-art segmentation algorithms into tracking pipelines. Nat Methods 19, 829–832 (2022). 10.1038/s41592-022-01507-1

