## Supplementary Figures for "Brain FGF2 and NCAM1 contribute to FGFR1-dependent progression of ER+ breast cancer brain metastases in young and aged hosts"

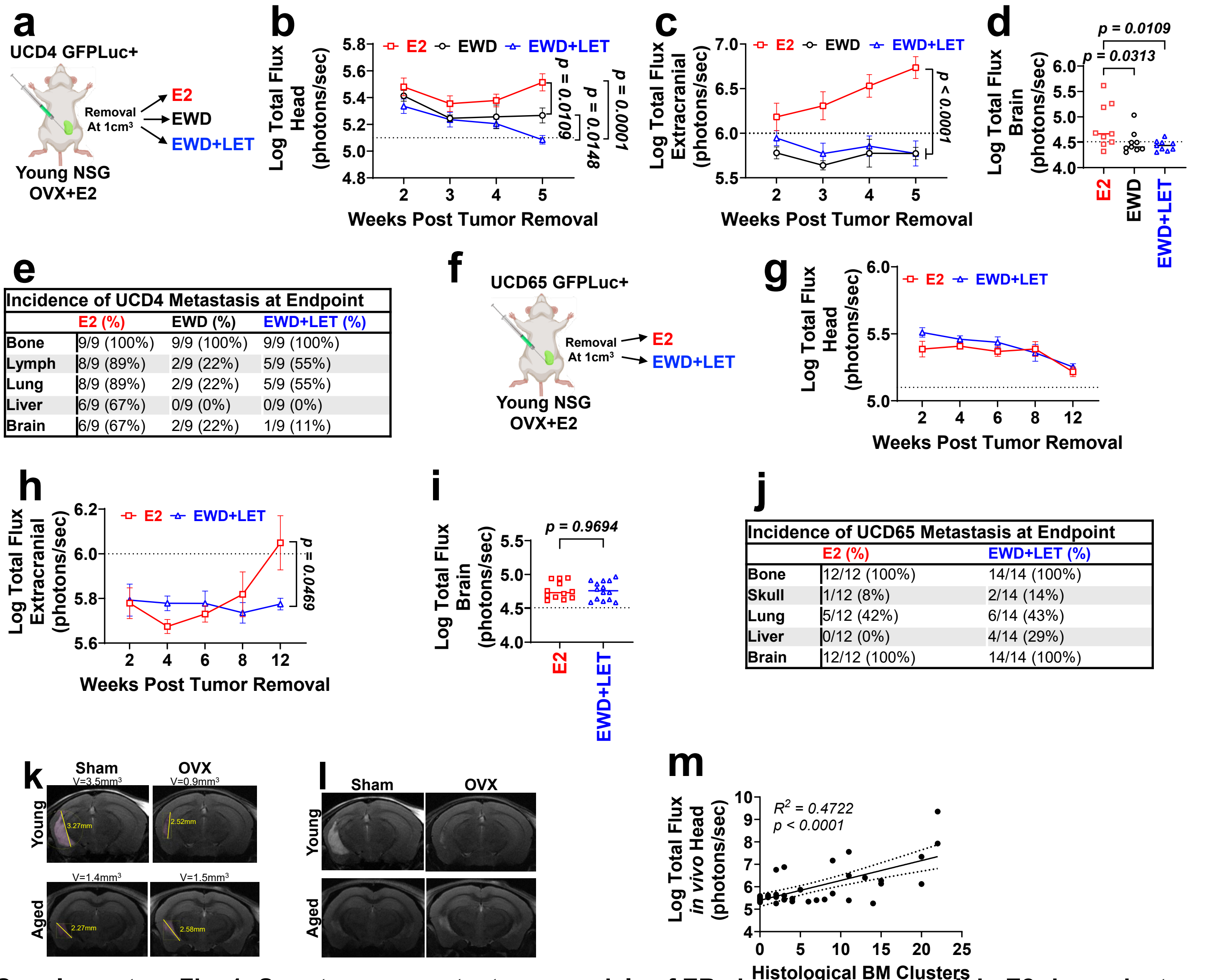


**e**

| Incidence of UCD4 Metastasis at Endpoint | | | |
|---|---|---|---|
| | E2 (%) | EWD (%) | EWD+LET (%) |
| Bone | 9/9 (100%) | 9/9 (100%) | 9/9 (100%) |
| Lymph | 8/9 (89%) | 2/9 (22%) | 5/9 (55%) |
| Lung | 8/9 (89%) | 2/9 (22%) | 5/9 (55%) |
| Liver | 6/9 (67%) | 0/9 (0%) | 0/9 (0%) |
| Brain | 6/9 (67%) | 2/9 (22%) | 1/9 (11%) |

**j**

| Incidence of UCD65 Metastasis at Endpoint | | |
|---|---|---|
| | E2 (%) | EWD+LET (%) |
| Bone | 12/12 (100%) | 14/14 (100%) |
| Skull | 1/12 (8%) | 2/14 (14%) |
| Lung | 5/12 (42%) | 6/14 (43%) |
| Liver | 0/12 (0%) | 4/14 (29%) |
| Brain | 12/12 (100%) | 14/14 (100%) |

**Supplementary Fig. 1: Spontaneous metastases models of ER+ breast cancer result in E2-dependent dissemination and outgrowth of metastases, as well as E2-independent metastatic progression in a subset of mice.**

**a,** UCD4 GFPLuc+ cells were injected into the mammary fat pad (MFP) of young NSG mice supplemented with pre-menopausal levels of E2. Tumors were removed at 1cm³ and mice were randomized to keep receiving E2 (n=9), undergo E2-withdrawl (EWD, n=9), or EWD+ letrozole (LET, n=9). **b,** Log-transformed head metastatic burden over time, measured by IVIS. **c,** Log-transformed overall metastatic dissemination over time, measured by IVIS. **d,** *Ex vivo* IVIS signal in the brain at endpoint. **e,** Quantification of metastasis incidence at multiple distant sites at endpoint. **f,** UCD65 GFPLuc+ cells were injected into the MFP of young NSG mice supplemented with E2. Tumors were removed at 1cm³ and mice were randomized to keep receiving E2 (n=12), or to undergo E2-withdrawl (EWD)+ letrozole (LET, n=14). **g,** Log-transformed head metastatic burden over time, measured by IVIS. **h,** Log-transformed extracranial metastatic dissemination over time, measured by IVIS. **i,** *Ex vivo* IVIS signal in the brain at endpoint. **j,** Quantification of metastasis frequency at multiple sites at endpoint. **k**, Masked T2-MRI images from **Fig. 1l** with maximum lesion diameter noted. **l**, Unmasked T2-MRI images from **Fig. 1l**. **m,** Correlation between IVIS head flux at endpoint and BM clusters via histological quantification. For **b,c,g,h** , lines denote mean ± SEM. Data were analyzed with 2-way ANOVA or mixed-effects analysis followed by Fisher's LSD test. P-value shown for last time point for all values <0.05. **d** was analyzed with one-way ANOVA followed by Fisher's LSD test. **i** was analyzed with an unpaired student's t-test. Gray dotted line denotes baseline IVIS signal from a tumor-free mouse.

**a**

| Cells: | UCD46 | UCD4 | UCD12 | UCD65 |
|---|---|---|---|---|
| % with BM | 0% | 20% | 40% | 80% |
| *FGFR* Copy # | No amp | 5x FGFR3 | 12x FGFR1 | 14x FGFR1 |
| *FGFR1* mRNA | 13.0 | 15.7 | 30.7 | 339.5 |
| *FGF1* mRNA | 0.00 | 0.00 | 0.00 | 0.31 |
| *FGF2* mRNA | 2.54 | 0.00 | 0.00 | 0.00 |
| *FGF4* mRNA | 0.00 | 0.00 | 0.00 | 0.00 |
| *FGF5* mRNA | 0.28 | 0.03 | 0.00 | 0.00 |
| *FGF6* mRNA | 0.00 | 0.00 | 0.00 | 0.00 |

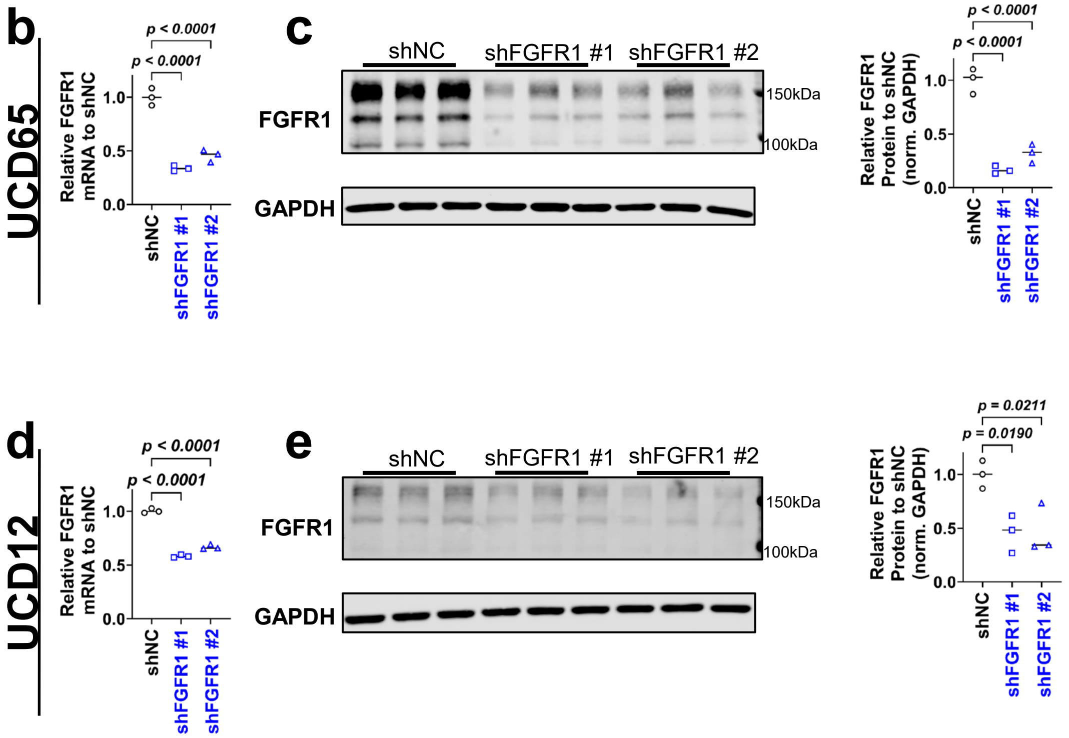


**Supplementary Fig. 2. FGFR1 is successfully downregulated in UCD65 and UCD12 cells.**
**a,** FGF mRNA expression (FPKM) in ER+ BC PDXs. **b,** RT-qPCR of FGFR1 mRNA in shNC and shFGFR1 UCD65 cells (n=3 replicates). **c,** Immunoblot of FGFR1 and quantification of FGFR1 normalized to GAPDH relative to shNC from 3 biological replicates of shNC and shFGFR1 UCD65 cells. **d,** RT-qPCR of FGFR1 mRNA in shNC and shFGFR1 UCD12 cells (n=3 replicates). **e,** Immunoblot of FGFR1 and quantification of FGFR1 normalized to GAPDH relative to shNC from 3 biological replicates of shNC and shFGFR1 UCD12 cells. Data in **b-e** were analyzed by 1-way ANOVA followed by Fisher's LSD test.

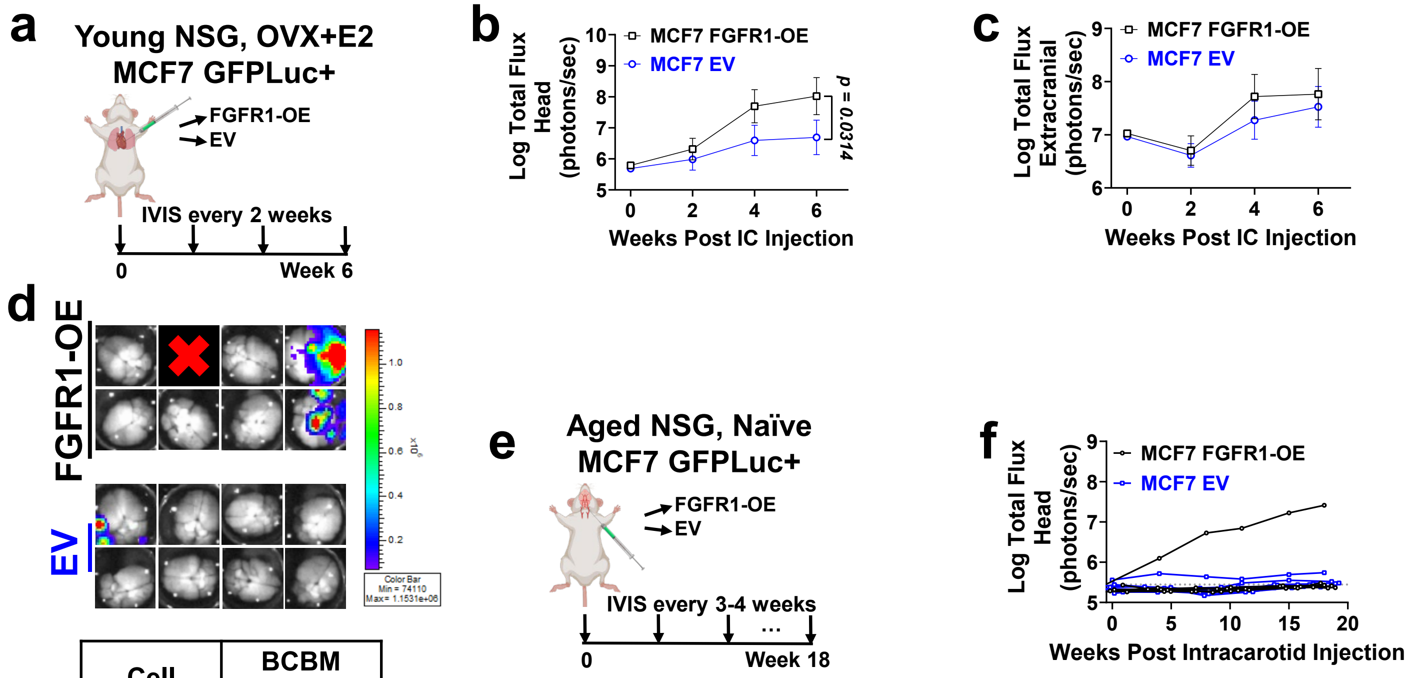


| Cell | BCBM Frequency |
|---|---|
| MCF7-EV | 1/8 (12.5%) |
| MCF7-FGFR1-OE | 3/8 (37.5%) |

**Supplementary Fig. 3: FGFR1 overexpression *in vivo*.**

**a,** FGFR1-OE (n=8) and EV (n=8) MCF7 GFPLuc+ cells were injected intracardially into young (<14w) NSG mice that were OVX+E2. **b,** Log-transformed head metastatic burden over time, measured by IVIS. **c,** Log-transformed extracranial metastatic burden over time, measured by IVIS. **d,** *Ex vivo* IVIS of brains, frequency of detectable BM. **e,** FGFR1-OE (n=7) and EV (n=9) MCF7 GFPLuc+ cells were injected via intracarotid artery into aged (>52w) naïve NSG mice. **f,** Log-transformed head metastatic burden over time, measured by IVIS. For **b**,**c**, lines denote mean ± SEM and data were analyzed with 2-way ANOVA followed by Fisher's LSD test.

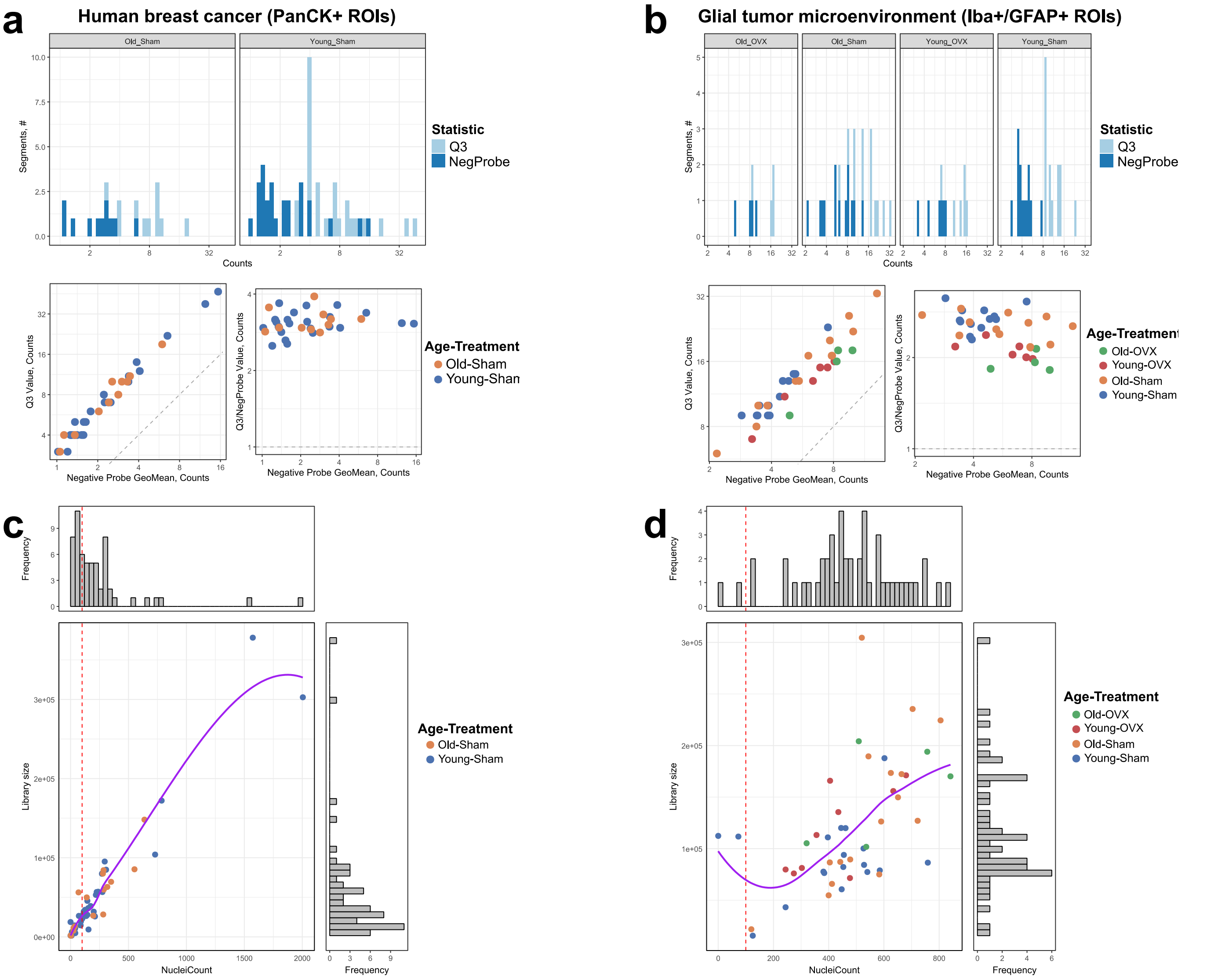


**Supplementary Fig. 4**: Quality control of NanoString libraries. DCC files were processed using the NanoStringNCTools, GeomxTools, and GeoMxWorkflows packages. Distribution of upper quartile (Q3) counts per segment (ROI) and of negative probe counts. Results for **a**, PanCK+ (human breast cancer) and **b**, Iba+/GFAP+ (glioma tumor microenvironment) ROIs. Distribution of library sizes and nuclei counts in the sequenced ROIs, excluding ROIs with a nuclei count below 100. Results for **c**, PanCK+ (human breast cancer) and **d**, Iba+/GFAP+ (glioma tumor microenvironment) ROIs.

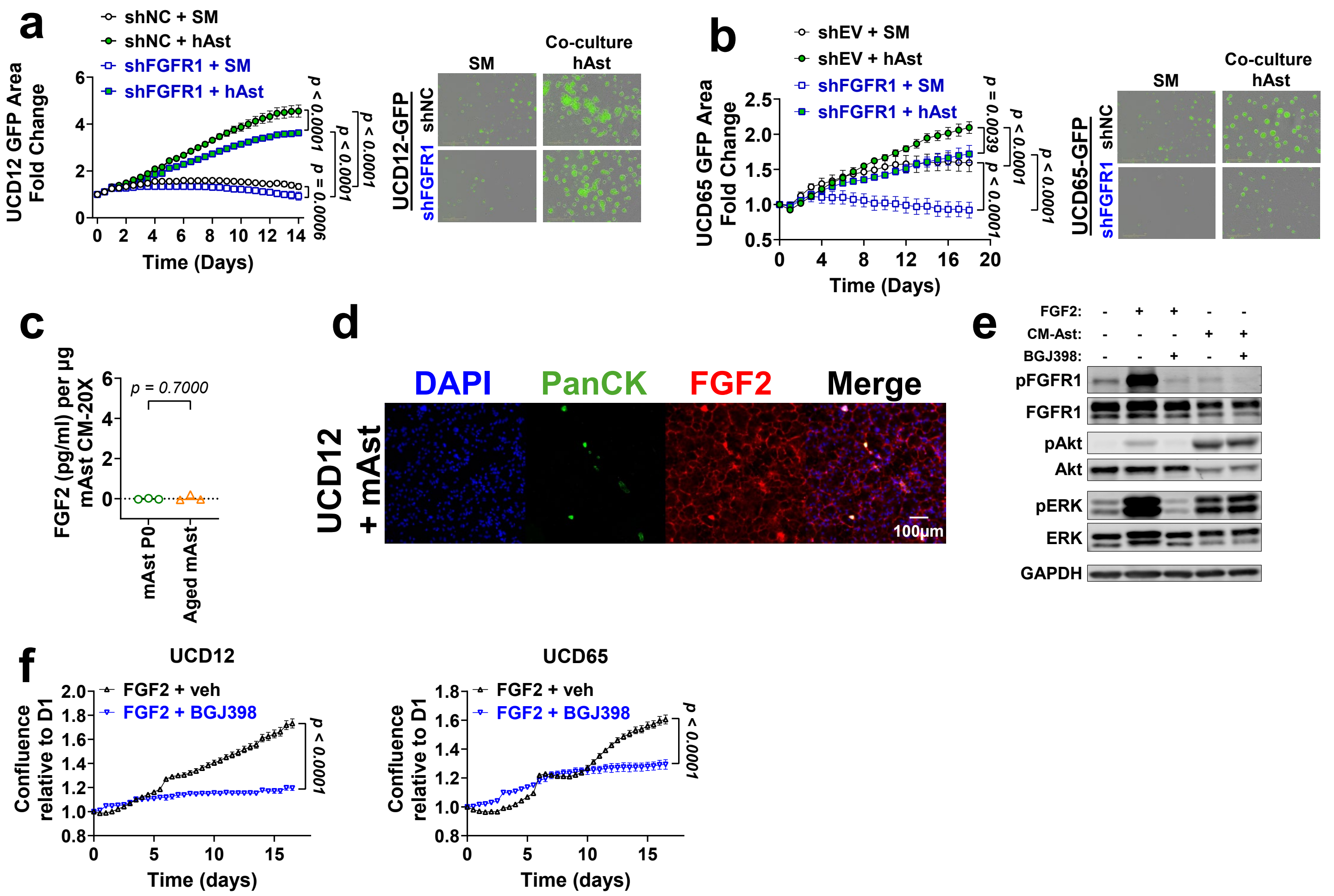


**Supplementary Fig. 5: Contribution of FGFR1 and human astrocyte co-culture to ER$^+$ cancer cell proliferation and tumor-initiating capability.**

shNC or shFGFR1 UCD12-GFP+ (**a**, n=3 replicates per condition) and shEV or shFGFR1 UCD65-GFP+ (**b**, n=5-6 replicates per condition) cells were treated with starvation medium (SM) alone or co-cultured with human astrocytes (hAst, ScienceCell #1800-10) in SM. Graphs show fold change in total green area relative to time 0, measured by Incucyte. Right: Representative images of UCD12 cells at 14 days and UCD65 cells at 18 days for each condition. **c,** FGF2 ELISA from 20X concentrated conditioned media from young and aged mouse primary astrocytes (n=3). **d,** IF staining of primary mAsts co-cultured with UCD12 cells stained for PanCK and FGF2. Scale bar: 100µm. **e,** Western blot of UCD65 cells starved for 72 hours, pre-treated with vehicle or BGJ398 (1µM, 1 hour), followed by FGF2 (10 ng/ml) or astrocyte-conditioned media (CM-Ast 10X) for 10min. **f,** Time course growth of UCD12 and UCD65 cells (n=6 replicates per condition) in E2-free media, treated with 10ng/mL FGF2 plus either vehicle or 10µM BGJ398. Data in **a,b,f** were analyzed with 2-way ANOVA followed by Fisher's LSD test. Data in **c** were analyzed by t-test.

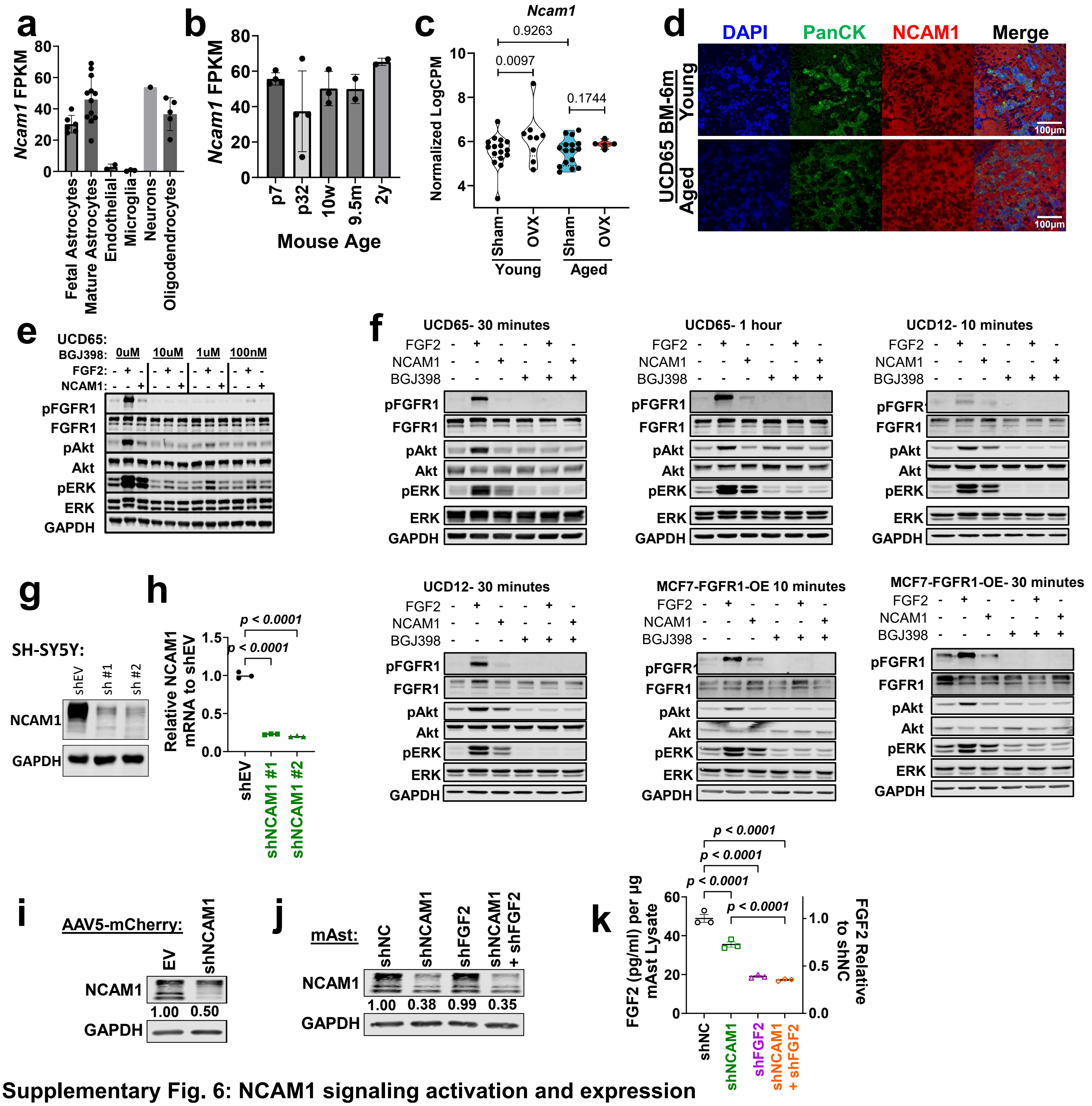


**Supplementary Fig. 6: NCAM1 signaling activation and expression**

**a,** Expression of NCAM1 mRNA in human brain cells as reported from brainrnaseq.org. **b,** NCAM1 mRNA expression in mouse astrocytes of different ages as reported at brainrnaseq.org. **c,** Normalized Log2 counts per million (CPM) for NCAM1 in non-glial ROIs from NanoString dataset (**Fig. 4f**), analyzed with empirical bayes. **d,** IF staining of PanCK and NCAM1 in brain metastases from UCD65 cells injected via intracarotid artery in young (<15w) and aged (>60w) NSG mice, 6 months post-injection. **e,** UCD65 cells were serum-starved and pretreated with 10µM, 1µM, or 100nM BGJ398 or vehicle for 1 hour, then treated with 10ng/mL FGF2, 250nM NCAM1, or vehicle for the indicated times. WBs of pFGFR1, FGFR1, pAkt, Akt, pERK, ERK, and GAPDH. **f,** UCD65, UCD12, and MCF7 FGFR1-OE cells were serum-starved and pretreated with 10µM BGJ398 or vehicle for 1 hour, then treated with 10ng/mL FGF2, 250nM NCAM1, or vehicle for the indicated times. WBs of pFGFR1, FGFR1, pAkt, Akt, pERK, ERK, and GAPDH. **g,** Immunoblot for NCAM1 in SH-SY5Y cells expressing shEV or shNCAM1. **h,** qPCR of NCAM1 in SY5Y shNCAM1 and shEV (n=3 replicates). 1-way ANOVA with Fisher's LSD test. **i,** Immunoblot for NCAM1 in mAsts infected with AAV5-mCherry EV or shNCAM1. **j,** Immunoblot for NCAM1 in mAsts expressing shNC, shNCAM1, shFGF2, or shNCAM1 + shFGF2. **k,** ELISA quantification of total and relative FGF2 protein in mAst lysates, analyzed by 1-way ANOVA with Fisher's LSD.

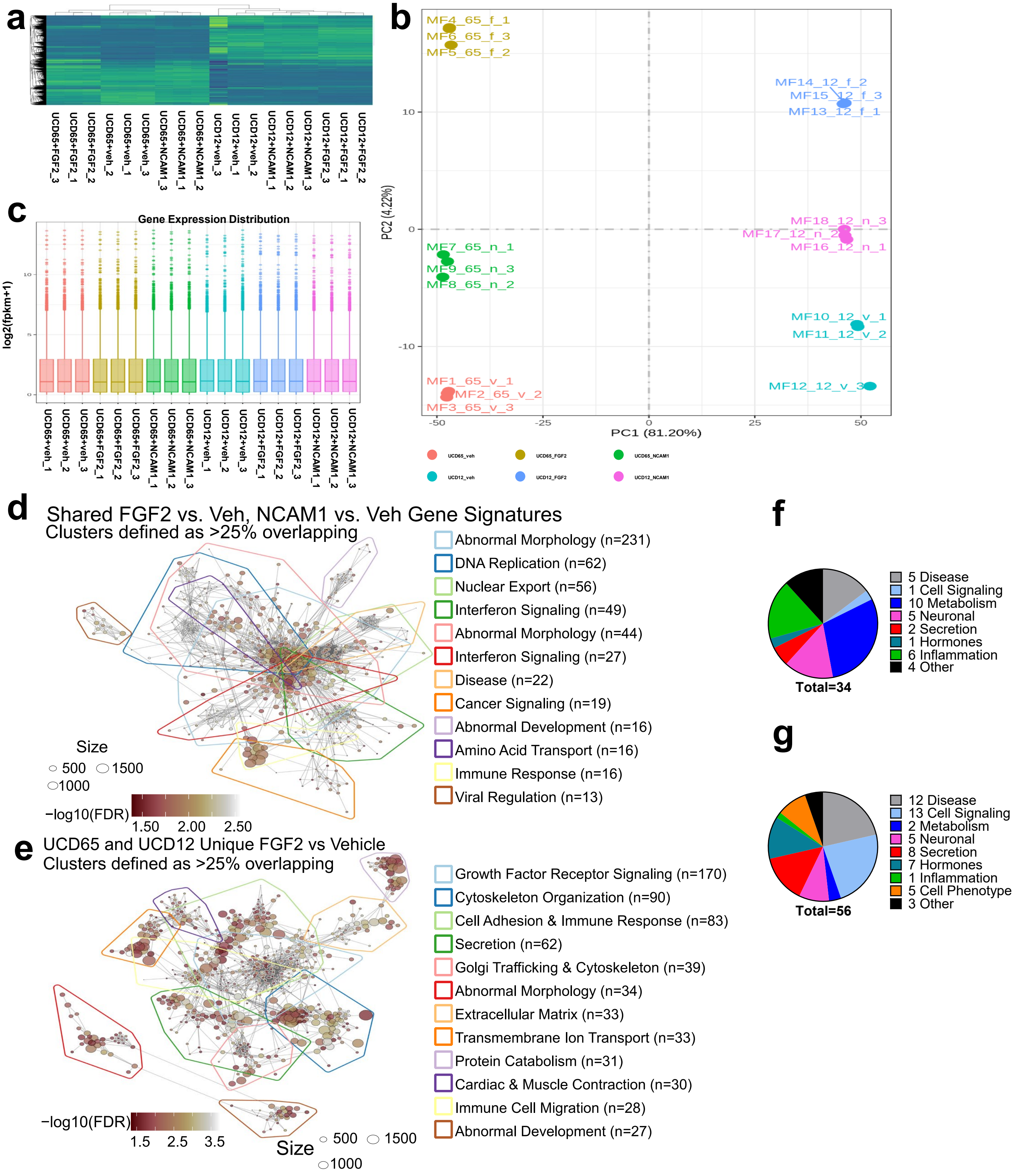


**Supplementary Fig. 7: RNA sequencing and analysis of NCAM1 and FGF2 treatments in UCD12 and UCD65 cells.**

**a,** Clustering of samples by normalized expression (z-score of normalized counts) demonstrates grouping by cell line and treatment. **b,** Principal component analysis (PCA) of gene expression values (FPKM) for each sample. **c,** Distribution of gene expression levels for each sample. **d,** Top 12 significant differentially expressed clusters of pathways (defined as signatures sharing >25% overlapping genes) for gene sets shared between FGF2_vs_veh and NCAM1_vs_veh. **e,** Top 12 significant differentially expressed clusters of pathways (defined as signatures sharing >25% overlapping genes) for gene sets unique to FGF2_vs_veh. **f,** Categorized gene signatures enriched following NCAM1 treatment. **g,** Categorized gene signatures repressed following NCAM1 treatment.

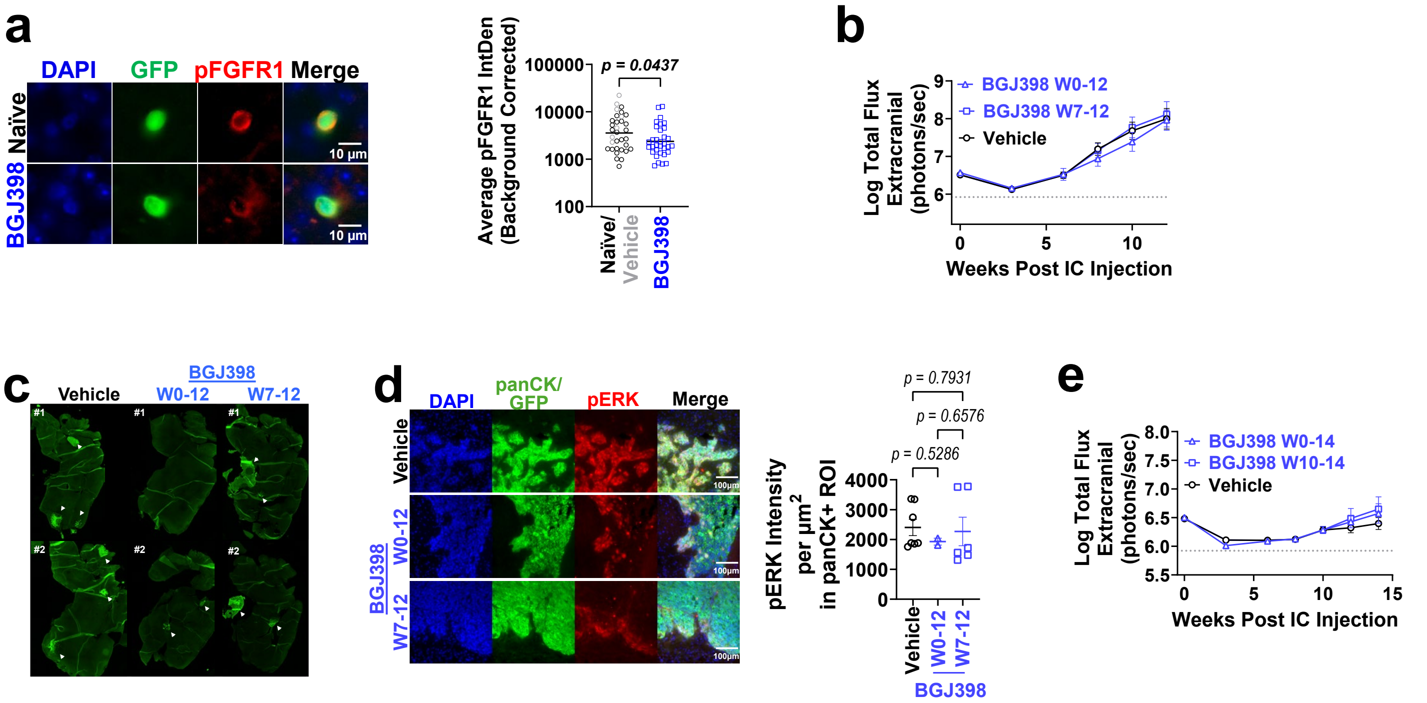


**Supplementary Fig. 8: FGFR inhibition *in vivo*.**

**a,** pFGFR1 quantification in brain disseminated UCD65 cells 5 days after intracardiac injection in NSG mice that were naïve or treated with BGJ398 5 days pre-injection. Graph shows average pFGFR1 intensity in GFP+ ROIs (n=33 images per group) corrected by background (GFP-). **b,** Log-transformed extracranial metastatic burden over time in young mice supplemented with E2, measured by IVIS. **c,** representative panCK IF stains of full sections of two animals per group. Macrometastases marked with white arrowhead. **d,** pERK expression and quantification in tissue from Fig. 8h. **e,** Log-transformed extracranial metastatic burden over time in aged naïve mice, measured by IVIS. For **a,** data were analyzed by lognormal t-test. For **d,** data were analyzed by 1-way ANOVA followed by Fisher's LSD. For **b,e,** lines denote mean ± SEM. Data were analyzed with 2-way ANOVA followed by Fisher's LSD test.
